# Temperature and food chain length, but not latitude, explain region-specific kelp forest responses to an unprecedented heatwave

**DOI:** 10.1101/2023.01.07.523109

**Authors:** Samuel Starko, Brian Timmer, Luba Reshitnyk, Matthew Csordas, Jennifer McHenry, Sarah Schroeder, Margot Hessing-Lewis, Maycira Costa, Amanda Zielinksi, Rob Zielinksi, Sarah Cook, Rob Underhill, Leanna Boyer, Christopher Fretwell, Jennifer Yakimishyn, William A. Heath, Christine Gruman, Julia K. Baum, Christopher J. Neufeld

**Affiliations:** Department of Biology, University of Victoria, 3800 Finnerty Rd., Victoria, British Columbia, V8P 5C2, Canada; School of Biological Sciences, University of Western Australia, Crawley 6009, WA, Australia; Bamfield Marine Sciences Centre, 100 Pachena Rd., Bamfield, British Columbia, V0R 1B0 Canada; Hakai Institute, Heriot Bay, BC V0P 1H0, Canada; Department of Geography, University of Victoria, 3800 Finnerty Road, Victoria, British Columbia V8P 5C2, Canada; Hornby Island Diving Ltd. 10795 Central Road Hornby Island BC V0R 1Z0 Canada; Coastal and Ocean Resources, 759A Vanalman Ave, Victoria, British Columbia V8Z 3B8 Canada; Mayne Island Conservancy, 455-A Dixon Road, Mayne Island, British Columbia V0N 2J2, Canada; Stqeeye’ Learning Society, PO Box 407 Salt Spring Island, British Columbia, V8K2W1, Canada; Pacific Rim National Park Reserve, Parks Canada Agency, 2040 Pacific Rim Highway, Ucluelet, British Columbia V0R 3A0, Canada; Project Watershed Society, 2536a Rosewall Crescent, Coutenay, British Columbia V9N 5N3, Canada; Huu-ay-aht First Nation, 170 Nookemus Road, Anacla British Columbia V0R 1B0, Canada

**Keywords:** climate change, extirpation, kelp forest, marine heatwaves, sea star wasting disease, trophic cascade

## Abstract

Kelp forests are among the most abundant coastal marine habitats but are vulnerable to the impacts of climate change. Between 2014 and 2016, an unprecedented heatwave and associated changes in trophic dynamics threatened kelp forests across the Northeast Pacific, with impacts documented from Mexico to Alaska. However, responses have varied substantially and remain poorly characterized across large stretches of coast, especially British Columbia (B.C.), which represents a significant percentage of the range of floating kelp species in the Northeast Pacific. Here, we explore variation in floating kelp (*Macrocystis, Nereocystis*) persistence pre- and post-heatwave across a >675 km latitudinal gradient, asking whether B.C. kelp forests are of conservation concern. We assembled and analyzed available quantitative kelp data, comparing snapshots of kelp extent before (1994 – 2007) and after (2018 – 2021) the heatwave in 11 regions spanning a range of temperature and sea otter-occupancy statuses, and contextualizing these with time series analyses, where available (n = 7 regions). We provide strong evidence that kelp forests have declined in many regions but with evidence of refugia at both local and regional scales. Kelp forest persistence was negatively correlated with summer sea temperatures in southern B.C., where temperatures varied by ~6°C across sites, at times exceeding species’ thermal tolerances. Kelp dynamics in northern regions appeared instead to be modulated by top-down control by urchins and sea otters. Our results demonstrate that B.C.’s kelp forest have been substantially reduced in recent years but that regional and local-scale factors influence the resilience of forests to large-scale perturbations.

## 1. Introduction

Human activities are altering the distribution and structure of marine ecosystems (Halpern et al., 2008; Steffen et al., 2011). Climate change, overfishing and pollution are among the drivers causing large-scale change in our ocean ecosystems (Jackson et al., 2001; Brierley & Kingsford, 2009; Mearns et al., 2010; Smale et al., 2019). In the face of these stressors, ecosystems at times undergo rapid regime shifts to states of less desirable structure and function (Scheffer & Carpenter, 2003). These shifts can result in the loss of habitat or productivity which may have cascading effects on organisms that use those ecosystems (Folke et al., 2004; Deyoung et al., 2008) and human communities that rely on them (e.g., Cesar, Burke & Pet-Soede, 2003; Pecl et al., 2017). Moreover, various ecological factors can reinforce regime shifts once they have occurred, potentially preventing ecosystems from returning to their initial state (Folke et al., 2004; Hughes et al., 2005; Filbee-Dexter & Scheibling, 2014; Filbee-Dexter & Wernberg, 2018), creating major challenges for conservation.

Coastal marine ecosystems can be especially sensitive to the impacts of climate change and other human activities with examples of regime shifts widespread across coral reefs (e.g., Graham et al., 2015; Arif et al., 2022), seagrass meadows (e.g., Moksnes et al., 2018; Chefaoui et al., 2021a) and seaweed communities (Filbee-Dexter & Scheibling, 2014; Filbee-Dexter & Wernberg, 2018). In temperate ecosystems, which often experience large seasonal and interannual fluxes in temperature and other climate-related variables, kelp forests are among the most abundant marine ecosystems (Jayathilake & Costello, 2021) but are threatened in many regions (Pörtner et al., 2019; Wernberg et al., 2019). Kelp forests provide essential habitat for a wide range of ecologically and economically important species, including fishes, invertebrates and other seaweed species (Steneck et al., 2002; Teagle et al., 2017; Shaffer, Munsch & Cordell, 2020). Moreover, they are highly productive and therefore fuel the growth of higher trophic levels (Duggins, Simenstad & Estes, 1989; Pessarrodona et al., 2022). Thus, declines in kelp forest abundance and extent can have far reaching consequences for nearshore ecosystems and beyond (Wernberg et al., 2019).

Evidence collected over the past two or more decades indicates that kelp forests are decreasing in abundance and extent across certain parts of the world due to combined effects of climate change and localized threats, including fishing, sewage run-off, invasive species and changes in freshwater outflow (Krumhansl et al., 2016; Filbee-Dexter & Wernberg, 2018; Wernberg et al., 2019; Hollarsmith et al., 2022). However, the trajectories of kelp forests around the world have been highly variable, with some regions showing stability (e.g., Chile and the Falkland Islands; Mora-Soto et al., 2021) or even increases in abundance (e.g., South Africa; Bolton et al., 2012), highlighting the importance of refugia at global, regional and local scales (Krumhansl et al., 2016; Wernberg et al., 2019). Where kelp forest ecosystem collapse has occurred, it has generally been associated with transitions to urchin barrens or communities formed by other (non-kelp) seaweeds (Wernberg et al., 2019), and there is evidence that transitions between these states can be challenging to reverse, often failing to return to the kelp forest state even after initial stressors are abated (Leinaas & Christie, 1996; Hughes et al., 2005; Pearse, 2006; Filbee-Dexter & Wernberg, 2018; Feehan, Grace & Narvaez, 2019). In some areas, kelp forest losses have had profound ecological and economic consequences from the collapse and closure of fisheries to detrimental impacts on tourism-based industries (Rogers-Bennett & Catton, 2019). In the USA, recent efforts have begun to determine whether one of the main kelp forest foundation species, bull kelp (*Nereocystis luetkeana*), should be listed and protected under the Endangered Species Act (Kelkar & Carden, 2022), highlighting growing efforts to restore and conserve kelp forest ecosystems in the face of ongoing climate change.

Between 2014 and 2016, a large-scale marine heatwave (MHW) known as “the Blob” unfolded along the west coast of North America (Lorenzo & Mantua, 2016; Tseng, Ding & Huang, 2017; Robinson, Yakimishyn & Evans, 2022), threatening kelp forests in many regions (Cavanaugh et al., 2019; Beas-Luna et al 2020; Starko et al. 2022). While warmer waters had direct impacts on kelp forests by imposing physiological stress and die-back (Cavanaugh et al., 2019; Starko et al., 2022), they also had indirect impacts on kelp forests by exacerbating the growing sea star wasting disease (SSWD) epidemic (Harvell et al., 2019; Hamilton et al., 2021). SSWD resulted in the functional extinction of *Pycnopodia helianthoides*, the sunflower star, across much of its distribution (Harvell et al., 2019; Hamilton et al., 2021), triggering trophic cascades that favoured sea urchins, the dominant herbivore of kelp forests (Schultz, Cloutier & Côté, 2016; Burt et al., 2018; Rogers-Bennett & Catton, 2019; McPherson et al., 2021; Starko et al., 2022). The combined effects of warming and expanding urchin populations have driven kelp forest losses throughout the Northeast Pacific (Beas-Luna et al., 2020) with severe impacts observed in populations of both major floating canopy-forming kelp species: giant kelp (*Macrocystis pyrifera*) and bull kelp (*Nereocystis luetkeana*).

Despite the growing evidence that MHWs and other extreme events have negatively impacted kelp forest ecosystems, the sheer extent and heterogeneity of these ecosystems makes it challenging to assess the scale of kelp deforestation. Kelp forests occupy more than one-third of the world’s coastlines, an area five times that of coral reefs (Jayathilake & Costello, 2021), and floating kelp occupy >30° latitude on the west coast of North America alone, suggesting that widespread declines could have profound impacts on the availability of coastal habitat for associated species and the extent of nearshore productivity. This could have important implications for economically critical fisheries that rely on kelp habitats, such as salmon and herring (Shaffer et al. 2019; Shaffer et al. 2020), throughout the Northeast Pacific and might also impact the extent to which coastal ecosystems draw-down and sequester carbon from the atmosphere at both global and regional scales (Krause-Jensen et al., 2018; Filbee-Dexter & Wernberg, 2020). To assess the spatial scale and extent of kelp forest loss, we must work to include historically understudied regions, including those that lack detailed multi-decadal time series, leveraging and scrutinizing all available data to draw inferences about kelp forest spatial contractions and associated concerns for conservation.

One region in which kelp forests have historically been understudied is British Columbia (B.C.), Canada. Past glaciation has left B.C.’s coast scarred with bays, fjords and channels that create inshore pockets of water that warm up in the summer to temperatures comparable to near the southern limit of either kelp species (Starko et al., 2022). For example, waters in both the Strait of Georgia and the west coast of Vancouver Island have reached temperatures greater than 20°C in recent summers (stations 3-5 in Fig 1; Starko et al. 2022). This is warmer than known growth optima for both canopy-forming kelp species (Supratya, Coleman & Martone, 2020; Fernández et al., 2020), suggesting that kelp forests may be threatened by these warm sea surface temperatures. Moreover, recent focal fieldwork along small stretches of the B.C. coast suggest that kelp forests have also declined in response to growing urchin populations (Schultz, Cloutier & Côté, 2016; Burt et al., 2018; Starko et al., 2022). However, the extent to which these threats are a concern across B.C.’s nearly 26,000 km coastline (more than twice that of California, Oregon and Washington combined) remains largely unclear.

**Fig 1.**
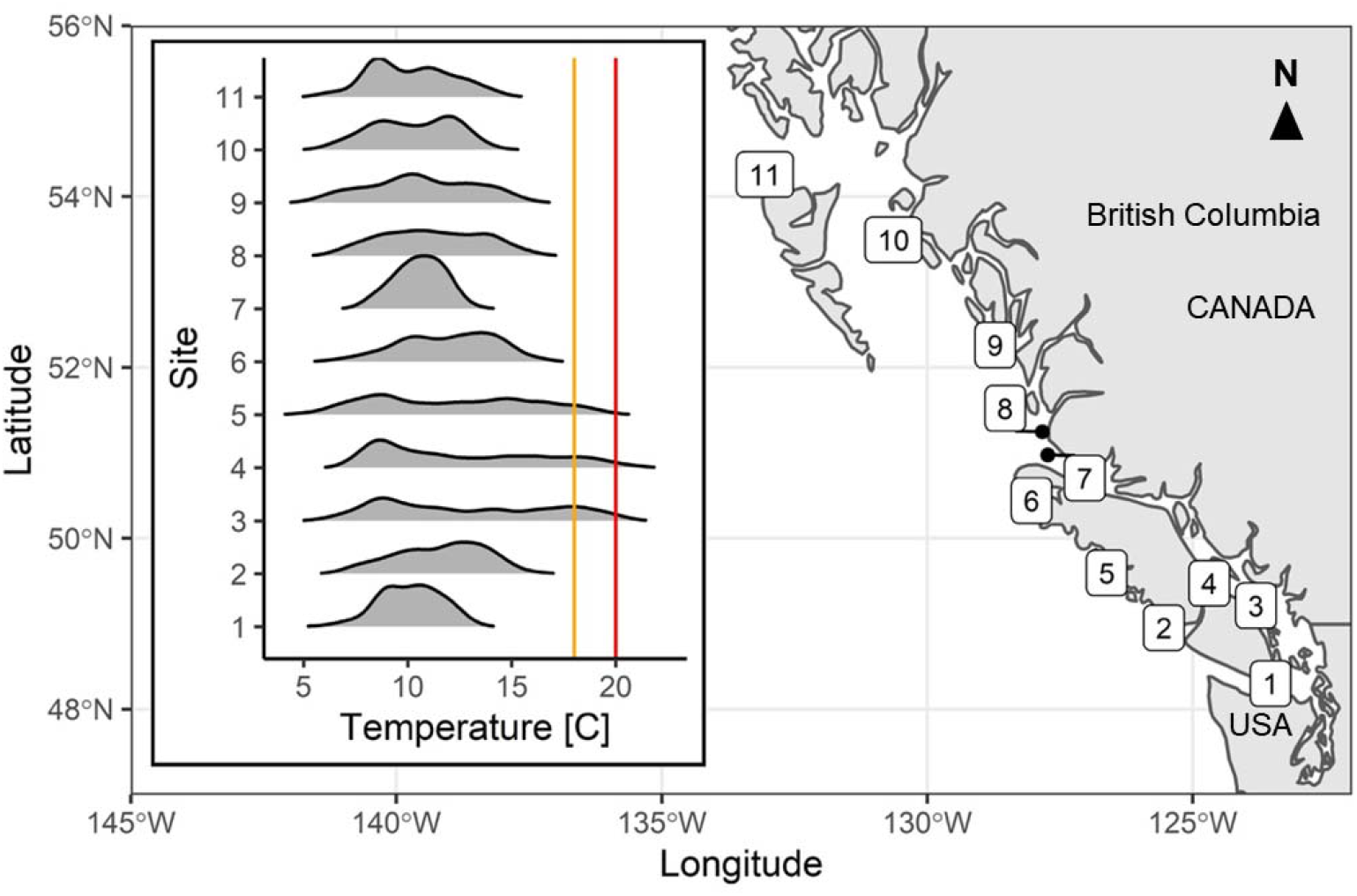
Sea surface temperature measured at high tide in the summers (June 21 – Sept 20) of 2014 to 2017. Data are from Fisheries and Oceans Canada Lighthouse Stations (labelled with numbers 1 – 11) along the coast of British Columbia. Inset plots indicate the relative density of all summer temperature measurements from 2014 to 2017. Note the lack of relationship between temperature and latitude. The orange line indicates 18°C and the red line indicates 20°C.

Here, we ask whether kelp forest extent in coastal B.C. has changed over the past two to three decades in response to recent environmental and biotic drivers. We assemble available quantitative data on kelp forest distributions from B.C. Because many regions lack long-term monitoring programs, we begin with numerous “snapshot” analyses–primarily using oblique shoreline photography supplemented with in situ data and satellite imagery when available–from two time points, one before recent heatwave and SSWD events (1994 – 2007) and one following (2017 – 2021). We then use time series data from all regions where they are available (n = 7) to contextualize large-scale patterns of change observed in the snapshot analyses. Moreover, we discuss how spatial patterns of change correlate with environmental variation and evidence of biotic interactions to make inferences about the drivers of kelp forest dynamics across a poorly studied region. This assessment aims to inform whether floating kelp forests (*Macrocystis* and *Nereocystis*) are of conservation concern in the province.

Various abiotic and biotic factors may make kelp more resilient in the face of warming and shifts in trophic dynamics (Hollarsmith et al., 2022). Spatial variation in temperature and other environmental variables can mediate the responses of kelp forests to large-scale warming (Wernberg et al., 2016; Cavanaugh et al., 2019; Starko et al., 2019; Filbee-Dexter et al., 2020), with factors like water motion, upwelling dynamics, and nutrient pollution leading to complexity in the distribution of environmental variation across the coastal fringe (Druehl, 1978; Hollarsmith et al., 2022; Spiecker & Menge, 2022; Starko et al., 2022). For example, while inland waters may warm up in the summer, areas on the outer coast with high wave exposure or currents can stay cool and nutrient rich through upwelling and mixing, potentially facilitating kelp persistence (Berry et al., 2021; Starko et al., 2022). On the other hand, food web length and structure can mediate the secondary impacts of SSWD. Specifically, when sea otters or other urchin predators are present they introduce functional redundancy, limiting the effects of sea star loss on the abundance of urchins and the subsequent effects on kelp forests (Burt et al., 2018; Eisaguirre et al., 2020). Consequently, here we test four hypotheses: 1) that kelp forests have experienced more losses (i.e., local-scale extirpations) than gains (i.e., local colonisations) across B.C.; 2) that kelp loss in some regions has been near-complete, similar to patterns observed in parts of California (e.g., Rogers-Bennett & Catton 2019); 3) that local environmental conditions (specifically temperature) have mediated the impacts of the 2014-2016 heatwave event on kelp forest distributions (with warmer areas more likely to experience kelp declines); and 4) that regions with sea otters have been more stable in the face of these large-scale perturbations due to top-down control on urchin populations.

## 2. Materials & Methods

### 2.1 Study design and regions

We selected study regions based primarily on the availability of imagery, and with the goals of capturing a range of sea surface temperatures as well as regions with and without sea otters. We were able to assemble data for 14 regions spanning a ~675-kilometer latitudinal gradient (6 regions for “snapshot analyses” only, 2 regions for time series only, and 5 regions for both types of analyses; see Tables 1–2). Our goal with the snapshot analyses was to quantify changes in kelp linear shoreline extent (hereafter, “extent”) using a timepoint from before (1995 – 2007) and a timepoint from after (2017 – 2021) the major 2014-2016 marine heatwave (MHW). While historical aerial images for snapshot analyses span more than a decade, all of these time points occurred before recent warming (since 2014) and SSWD that were known to drive major changes to kelp forest dynamics in the Northeast Pacific (Beas-Luna et al., 2020). We also compiled various time series data where available; while datasets differ in methodology and response variable (e.g., linear extent vs total kelp area; see below), they were internally consistent and therefore provide important context for patterns of change captured by snapshot analyses.

**Table 1.** Study regions used to assess changes in canopy kelp distribution from “snapshot” analyses. Asterisk (*) indicates regions that were also included in time-series analyses. For kelp species present: MP = *Macrocystis pyrifera*, NL = *Nereocystis luetkeana*, † indicates which species is more common when both are present. For otter status: + = increasing populations, - = otter populations not present.

| Region | Region Name | Latitude | Longitude | Year prior to MHW | Year following MHW | Otter status | Kelp species present |
| --- | --- | --- | --- | --- | --- | --- | --- |
| i | Valdes and Gabriola Islands | 49.0 to 49.2 | -123.6 to -123.8 | July 29, 2004 | August 7, 2021 | - | NL |
| ii* | Mayne and Saturna Island | 48.8 to 48.9 | -123.0 to -123.4 | July 29, 2004 | August 7, 2021 | - | NL |
| iii* | Cowichan Bay | 48.7 to 48.8 | -123.5 to -123.7 | September 24, 2004 | July 27, 2017 | - | NL |
| iv | Juan de Fuca Entrance | 48.5 to 48.6 | -124.3 to -124.7 | August, 13-14, 2007 | August 9, 2021 | - | NL |
| v | West Coast Trail | 48.6 to 48.8 | -125.2 to -125.3 | August 14, 2007 | August 8-9, 2021 | - | MP, NL† |
| vi* | Barkley Sound | 48.8 to 48.9 | -125.0 to -125.2 | August 14, 2007 | August, 2018 | - | MP†, NL |
| vii | Nootka Sound | 49.4 to 49.6 | -126.5 to -126.8 | June 26, 1994 | July 24, 2021 | + | MP†, NL |
| viii | Quatsino Sound | 50.3 to 50.5 | -127.5 to -128.2 | May 17, 1999 | June 17, 2018 | + | MP†, NL |
| ix* | South Central Coast* | 51.8 to 52.2 | -128.2 to -128.6 | July 21, 1997 | May 18, 2018 | + | MP†, NL |
| x* | Laredo Sound* | 52.4 to 52.8 | -128.8 to -129.3 | July 24, 1997 and July 12-13, 1998 | July 7, 2019 | - | MP, NL† |
| xi | Dundas Island | 52.6 to 54.4 | -130.7 to -131.0 | July 2, 2000 | July 4, 2019 | - | MP, NL† |

**Table 2.** Summary of data sources used for time series analysis. Asterisk (*) indicates regions that are also included in the snapshot analyses. Note that time series are generally not continuous but include some gap years. For specific years and dates used in each time series, see Table S3.

| Region | Region Name | Start date | End date | Number of time points | Data type | Methods |
| --- | --- | --- | --- | --- | --- | --- |
| **i-b** | Central Strait of Georgia | 2013/2014 | 2022 | n = 24-34 per site | Presence-absence at two sites | SCUBA surveys and videos |
| **ii** | Mayne Island* | 2010 | 2021 | n = 8 | Kelp forest area (m<sup>2</sup>) | Kayak surveys |
| **iii** | Cowichan Bay* | 2004 | 2017 | n = 5 | Presence-absence of shoreline segments (% occupied) | High resolution satellite imagery |
| **vi** | Barkley Sound* | 2007 | 2022 | n = 6 | Presence-absence of shoreline segments (% occupied) | Aerial image, satellite, boat surveys |
| **ix** | South Central Coast* | 1984 | 2021 | n = 38 | Kelp forest area (m<sup>2</sup>) | LandSat satellite imagery |
| **ix-b** | Calvert Island | 2008 | 2022 | n = 7 - 10 per site | Kelp forest area (m<sup>2</sup>) at two sites | RPAS |
| **x** | Laredo Sound* | 2007 | 2019 | n = 3 | Presence-absence of shoreline segments (% occupied) | Aerial image, satellite |

### 2.2 Snapshot analyses of kelp linear extent

To assess changes in linear extent (measured here as presence-absence of kelp along shoreline segments), we performed analyses focused on two time points: one before the MHW and one after (hereafter “snapshot analyses”). For nine regions, we used oblique aerial imagery collected by the ShoreZone initiative (Howes, Harper & Owens, 1994; Cook et al., 2017) and Environment & Climate Change Canada (ECCC) between 1995 and 2021 as data sources for both time points. For one of the regions (Barkley Sound), more recent ShoreZone imagery from before the MHW were available (2007) and we coupled these with in situ surveys conducted in 2018, while for another region (Cowichan Bay), we used two years from a dataset derived from high resolution satellite imagery (see below). A summary of data sources for each region is provided in Table S1

For the ten regions that involved oblique aerial imagery (including Barkley Sound), we created shoreline segments to classify stretches of shoreline that could be identified in both pre- and post-MHW imagery, and where either one or both of the images contained kelp canopy that was clearly visible in oblique images. Oblique imagery was taken at low tidal heights when most kelp canopy can be expected to be floating at the surface (Schroeder et al., 2019; Timmer et al., 2022), but since the imagery was collected at an oblique angle, we were unable to accurately assess changes in the area of kelp canopy over time and restricted these analyses to presenceabsence. Therefore, kelp canopy was determined to be either present or absent within each image for each segment, and the segment was accordingly classified as either a ‘gain’ (colonisation; absent in pre-MHW imagery but present in post-MHW imagery), a ‘loss’ (extirpation; present pre-MHW but absent after), or as ‘stable’ (kelp remained present at both time points) for each segment between the two time periods. Out of an abundance of caution, if the kelp in either image was not clearly identifiable due to glint on the water surface, choppy water, or if the image was too grainy to reliably identify kelp, the segment was not used. For these ten regions, the methods varied slightly depending on the availability of imagery and/or the length of shoreline surveyed.

In six regions (Valdes/Gabriola, Nootka, Quatsino, South Central Coast, Laredo Sound, Dundas Island), coverage of historical images was limited. For these regions, shoreline segments (~30-60 m in length) were established based on recognizable shoreline features in the aerial images that could be georeferenced in google earth. Due to data limitations in these regions, some segments were created based on stills taken from oblique-facing aerial videos that were collected during initial surveys rather than images themselves. Most photos (from either video stills or original photography) only covered a single segment, but in some cases, two or three segments were established from wider shot photography. To capture colonisation events or determine the extent to which some stretches of shoreline lacked kelp in both survey years, we also collected observations of kelp absences for any images where kelp was not present in the historical year and for which there was modern imagery of that same stretch of coastline that also lacked kelp. We did not quantify the length of shoreline associated with absence observations unless they had been colonised by kelp between the time points. In this latter case, shoreline segments were created as described above to capture kelp colonisation of a new stretch of shoreline.

For three regions, where historical oblique imagery was not limiting and virtually the entire coast was photographed at high quality (Juan de Fuca Entrance - region iv, West Coast Trail – region v, Barkley Sound – region vi), different approaches were employed to systematically survey photographic data. For the West Coast Trail (region v) and Juan de Fuca entrance (region iv), which collectively represent more than 80km of shoreline, a systematic subsampling method was used. First the coastline was split into 250m pixels, and at the start of each pixel, a single shoreline segment (length ~20-100m depending on shoreline features; West Coast Trail average = 48m; Juan de Fuca average = 40m) was established, provided the image met the quality criteria described above in imagery from both time points. For Barkley Sound (region vi), we used two time points from a time series involving multiple data sources (see Starko et al 2022 and description below). In short, we systematically resurveyed a 16km stretch of coast (originally surveyed in 2007) in 2018, 2021 and 2022 (see Starko et al. 2022). The coast was segmented into ~20 – 100m segments (32m on average) segments as above. For the snapshot analysis, we compared data from 2007 to data from 2018.

Shoreline segments (100m in length) in Cowichan Bay (region iii) were established as described in Schroeder et al. (2020) and relied on high resolution satellite imagery (Digital Globe; 2.5m resolution or higher) rather than oblique aerial imagery. For this dataset, a time series was produced from imagery in 2004, 2012, 2015, 2016 and 2017 from <2m tidal elevation in July – September, from which we used data from 2004 and 2017 for the snapshot analysis. Methodological details for this dataset are given in section 2.3

### 2.3 Time series analysis

We assembled time series datasets for 7 regions, five of which directly overlap with regions examined in the snapshot analyses (see Table 2 for summary of years and data types). These five regions are Mayne Island (region ii), Cowichan Bay (iii), Barkley Sound (vi), South Central Coast (ix) and Laredo Sound (x). In contrast, the Central Strait of Georgia region (region i-b), which includes a stretch of the Strait of Georgia from Denman Island down to Nanaimo is adjacent to the Valdes/Gabriola region (region i) but does not include the same stretches of shoreline (see Fig 2, Fig S1). Similarly, Calvert Island (region ix-b) is adjacent to the South Central Coast (region ix) but is not contained within the region analysed for the snapshot analysis (see Fig 2, Fig S2). Data from Central Strait of Georgia (region i-b) involved analysis of SCUBA diver observations from 2 sites (see below), and Calvert Island (region ix-b), involved the analysis of aerial imagery acquired from remotely piloted aerial systems (RPAS) flown at two sites.

**Fig 2.**
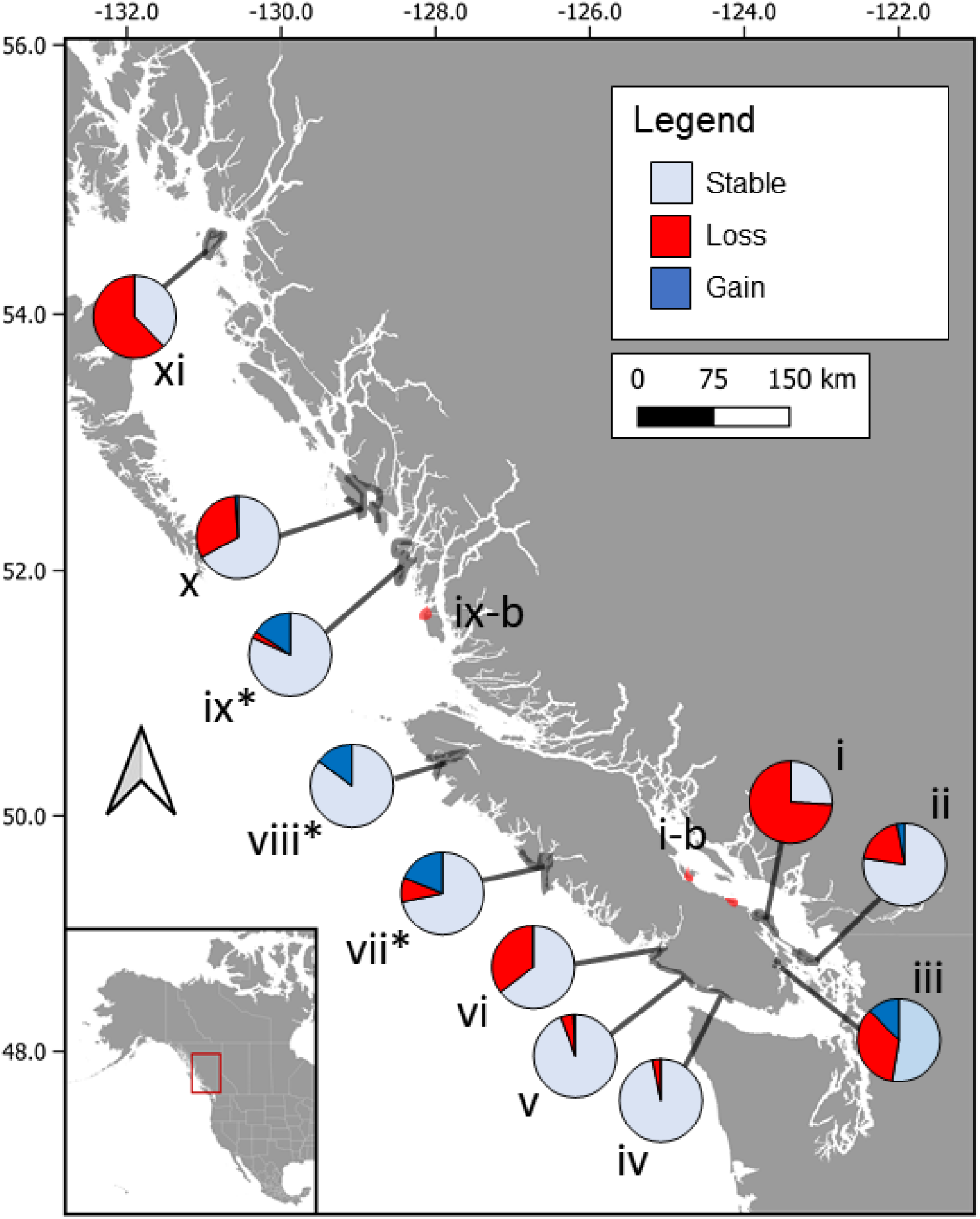
Regional-scale patterns of kelp persistence comparing pre-(1997 – 2007) and post-marine heatwave (post-2016) imagery. Pie charts indicate the relative direction of change in canopy kelp presence (both *Macrocystis* and *Nereocystis*), at the level of individual segments, between the two time points based on shoreline segments visible in imagery from both time points. The regions with expanding otter populations are indicated with asterisks. Regions are as follows: i) Valdes/Gabriola, ii) Mayne/Saturna, iii) Cowichan Bay, iv) Juan de Fuca Entrance, v) West Coast Trail, vi) Barkley Sound, vii) Nootka Sound, viii) Quatsino Sound, ix) South Central Coast, x) Laredo Sound, xi) Dundas Island. Region only included in the time series analysis are shown in red: i-b) Central Strait of Georgia, ix-b) Calvert Island.

Where available, in situ data were used for time series analyses. This was the case for two regions: the Central Strait of Georgia (region i-b) and Mayne Island (ii). To contextualize changes in bull kelp (*Nereocystis*) occupancy (i.e., presence-absence) in the Central Strait of Georgia (where *Macrocystis* is not present), we extracted data from citizen SCUBA diver videos (available online or through a local dive shop) and from the logbooks of authors. At one site, Eagle Rock (Denman Island), dives were conducted intentionally looking for kelp forests by three of the authors (A. & R. Zielinski, W. Heath) as part of a restoration and monitoring initiative. Here, logbooks recorded whether *Nereocystis* was present and this was converted into presence-absence data for the site. At the other site (Tyee Cove, Nanoose Bay), we assembled recreational SCUBA diver videos (from a local dive shop and online – e.g., YouTube; Table S2) and determined whether *Nereocystis* was present in each video over an 11-year period. While videos were generally not taken with the intention of tracking kelp, *Nereocystis* was a frequent occurrence in the shallows of this site alongside other kelp species, and therefore we would expect it to appear in citizen SCUBA diver videos either on purpose or incidentally while filming in the “kelp zone”. We included observations from April – October, to reflect the growing season of *Nereocystis*. While we included all videos or blog posts where *Nereocystis* was visible as observations of canopy kelp being “present”, we required that videos cover at least 20 seconds of footage in the “kelp zone” (i.e., understory kelp present), where conditions would be suitable for bull kelp, to include a video as an absence observation. Both sites were selected based on direct observations by the authors (Timmer, Heath, A & R. Zielinski) that the forests had disappeared in recent years and therefore may offer insight into the timing of losses but not the extent of losses, nor does this capture interannual variation in abundance or extent of kelp forests that have persisted in this region.

For the second time series derived from in situ data (Mayne Island – region ii), data from citizenscience kayak surveys were analyzed to produce a time series. Surveys were conducted in situ by encircling the surface extent of kelp forests during low tides <1.2m above chart datum and taking GPS points to identify the perimeter of the bed. Polygons were then produced from these data to represent kelp extent at each time point. Because survey areas varied in their spatial coverage between years, polygons were clipped according to spatial overlap of the survey areas to maximize the temporal coverage of the surveys. The resulting dataset covered five discontinuous sites (Fig S1) that were each sampled 9 times between 2010 and 2022.

For five other regions (iii, iv, ix, ix-b, x), various approaches were used to construct time series with remote sensing technologies (satellites, aerial images from piloted vehicles, RPAS), with analysis type dependent on data availability and/or previously conducted region-specific analyses. While the Barkley Sound (region vi) time series involved a combination of remotely sensed and in situ data, all other regions used remote sensing data for all time points. A summary of remote sensing data sources used for time series is provided in Table S3.

For Cowichan Bay (region iii), high resolution WorldView-2 satellite images were acquired at tidal height below 2.0 m from July, August and September, corresponding to the growing season bull kelp (see Tables 2, S3 for years). In short, kelp presence and absence along each shoreline segment was assessed using an unsupervised ISODATA classification approach, considering land and 30 m bathymetry masks and a buffer along the shoreline to minimize the effects of adjacency. This data set is published and presented by Schroeder et al. (2020).

For Barkley Sound (region iv), aerial and high resolution satellite images were classified visually (see Starko et al., 2022) and compared to boat surveys conducted between 2018 and 2022 (see snapshot methods above). The same shoreline segments were used as described in section 4.2, however the dataset was trimmed to ensure only the subset of segments present in all years of the time series were analysed.

For the South Central Coast (ix), the Google Earth Engine Kelp Mapping Tool was used to produce a time series of annual maximum kelp extent from 1984 to 2021 using methods described within Nijland et al. (2019). In brief, a time series of maximum annual kelp area (m^2^) was derived using the Normalized Difference Vegetation Index (NDVI) from Landsat 5 TM, 7 ETM+ and 8 OLI imagery with a minimum NDVI threshold of 0.02 and detection threshold of 2 (ie. each pixel had to be detected above the minimum NDVI threshold twice in each time period to be classified as kelp) for all image scenes available between May 1 – Oct 31. In some years, detection thresholds were changed due to limitations in available imagery (mostly due to cloud cover). Images used had cloud cover < 90% and a tidal stage of <3.5 m (chart datum) of each year. Where the Landsat cloud mask was found to perform poorly, some image scenes were removed manually. A land mask was applied with a 30 m buffer (1 Landsat pixel) to remove potential mixed pixels containing land which could be falsely detected as kelp. Therefore, these data outputs consider “offshore” kelp explicitly.

For Calvert Island (ix-b), total canopy kelp area was quantified at two sites using imagery from RPAS flown in situ. Meay Channel is a site with *Macrocystis*, while North Beach is a *Nereocystis* site. Total kelp area at each time point was assessed either manually or using an index and threshold method.

For Laredo Sound (x), aerial imagery (from 2007), visible colour satellite imagery from 2013 (Google Earth) and oblique imagery from ECCC (2019) were compared and classified visually (as with the Barkley Sound time series) using 50m segments (presence-absence only).

### 2.4 Environmental data

We used environmental data to test 1) whether spatial patterns of temperature predicted kelp persistence based on snapshot analyses; and 2) how kelp area in the longest time series (South Central Coast) correlates with temporal temperature anomalies. To assess how patterns of kelp persistence in snapshot analyses relate to local summer sea surface temperatures, we used average daily sea surface temperature from the LiveOcean Model, a Regional Ocean Modeling System adapted to the coastal waters off of Washington, Oregon, and southern British Columbia (Fatland, MacCready & Oscar, 2016). We extracted and averaged data from August 2017 (the first year of the model) which, although not during the 2014-2016 MHW, was still an anomalously warm year. This model has a grid size of 500 – 1500m (depending on location) which may miss some fine-scale temperature variation. It captures known temperature gradients on southern Vancouver Island such as in Barkley Sound (Starko et al. 2022) and the Salish Sea (Ban et al., 2016) but does not include the three regions north of Vancouver Island. To assess temperature anomalies relevant to the South Central Coast (region ix) time series, we used a temperature time series from McInnes Lighthouse which is nearby this region. We then calculated average month anomalies using data from 1982 to 2012 as the baseline.

### 2.7 Sea otter occupancy status

We used previously published reports (Nichol et al., 2015, 2020) to infer which of the 14 regions included in this study are occupied by sea otters (*Enhydra lutris*). These reports document surveys conducted to quantify population size and distribution of otter populations across the coast of B.C. These results show that otters are consistently present (and with growing populations) in three regions examined here: Nootka Sound (vii), Quatsino Sound (viii) and the South Central Coast (ix). Although considered to have expanded to areas around Calvert Island (region ix-b) in 2013 (Nichol et al., 2015), on-the-ground observations and surveys have shown the occupation of focal sites (North Beach and Meay Channel) in this region was short-lived and otters were no longer using these sites after 2016 despite being present on nearby islands.

### 2.6 Statistical analysis

To test whether regions varied in their trajectories in the snapshot analysis, we used a Fisher’s exact test to determine whether kelp change status (stable, gain, loss) was contingent on region. We also tested whether summer sea surface temperature (using modelled data from August 2017) predicted kelp persistence from snapshot data in southern regions using a spatially explicit binomial glm (0 = kelp loss, 1 = kelp gain or persistence). We then used various statistical models to determine whether kelp forest extent or abundance has changed through time in the time series data. Specifically, we used linear models to test for change through time for Barkley Sound (extent), Cowichan Bay (extent) and Mayne Island (kelp forest area) regions because data were continuous and normally distributed. For Laredo Sound, we tested whether years differed in their linear extent using a binary glmm (fixed = year [categorical]; random = segment). For Calvert Island region, where data were continuous but non-normal, we tested for change through time across both sites using a glmm fit with a gamma distribution (fixed = year [continuous]; random = site). Finally, we tested for change in offshore kelp forest area through time in the South Central Coast time series using a linear model and also tested whether kelp forest area following 2014 was drawn from the same distribution as prior to the heatwave using a Wilcoxin rank sum test.

## 3. Results

We found substantial changes in the linear extent of kelp forests when comparing pre- and post-MHW data from snapshot analyses; however, the direction and amount of change varied across regions (Fig 2; Fisher’s Exact Test: p = 0.0005). Of the 11 regions that we examined with our snapshot analyses, 6 had more kelp losses than gains, 2 had more gains than losses, and 3 had roughly no change (<10% net loss or gain). Regions that experienced the greatest kelp loss spanned the entire latitudinal gradient from Valdes/Gabriola – region ii (74 % loss; ~49° N) to Laredo Sound – region x (31% loss; ~53°N) and Dundas Island – region xi (62% loss; ~55°N). Moreover, losses were observed in regions dominated by both *Macrocystis* and *Nereocystis* (e.g., Barkley Sound – region vi versus Valdes/Gabriola – region i). In contrast, the South Central Coast (region ix) and Quatsino Sound (region viii) regions experienced increases in linear extent compared to historical snapshots, with 16% and 15% gains, while only experiencing 3% and no losses, respectively. Regions on the exposed outer coast of southern Vancouver Island (Juan de Fuca – region iv, West Coast Trail – region v) experienced very little change in kelp extent (<5% net change) between the two time points while Nootka – region vii – experienced both increases and decreases, resulting in a net change of only ~9% (Fig 1). Regions that experienced increases or stability also included those dominated by both canopy kelp species.

In addition to differences amongst regions, spatial patterns in kelp persistence tended to reflect the mediating impacts of fine-scale environmental variation. In southern regions without otters (regions i to vi), kelp loss strongly correlated with local summer sea surface temperatures. Coastlines along the southeast (regions i to iii) and southwest sides (regions iv to vi) of Vancouver Island span multiple local and regional temperature gradients (see Fig 3) and kelp persistence patterns from these regions strongly correlate with this fine-scale variation in temperature. Kelp loss on Mayne/Saturna (region ii) was largely restricted to the northeastern sides of islands that experience greater temperatures than other parts of the region (Fig 3). Similarly, the Valdes/Gabriola regions, which experience particularly warm summer temperatures, experienced significant losses across the entire region. In Barkley Sound, kelp forests disappeared primarily from inner parts of the region where conditions are known to get much warmer (see Starko et al., 2022; Fig 3) while remaining towards the outer shore, including adjacent outer shore regions (West Coast Trail, Juan de Fuca Entrance). We specifically tested whether spatial variation in temperature predicted kelp forest persistence within and across all regions on southern Vancouver Island (n = 6 regions; n = 798 segments) and found that, after accounting for spatial autocorrelation, kelp forest persistence was strongly predicted by local temperature variation (Spatial GLM: *X*^2^ = 24.402, p < 0.0001).

**Fig 3.**
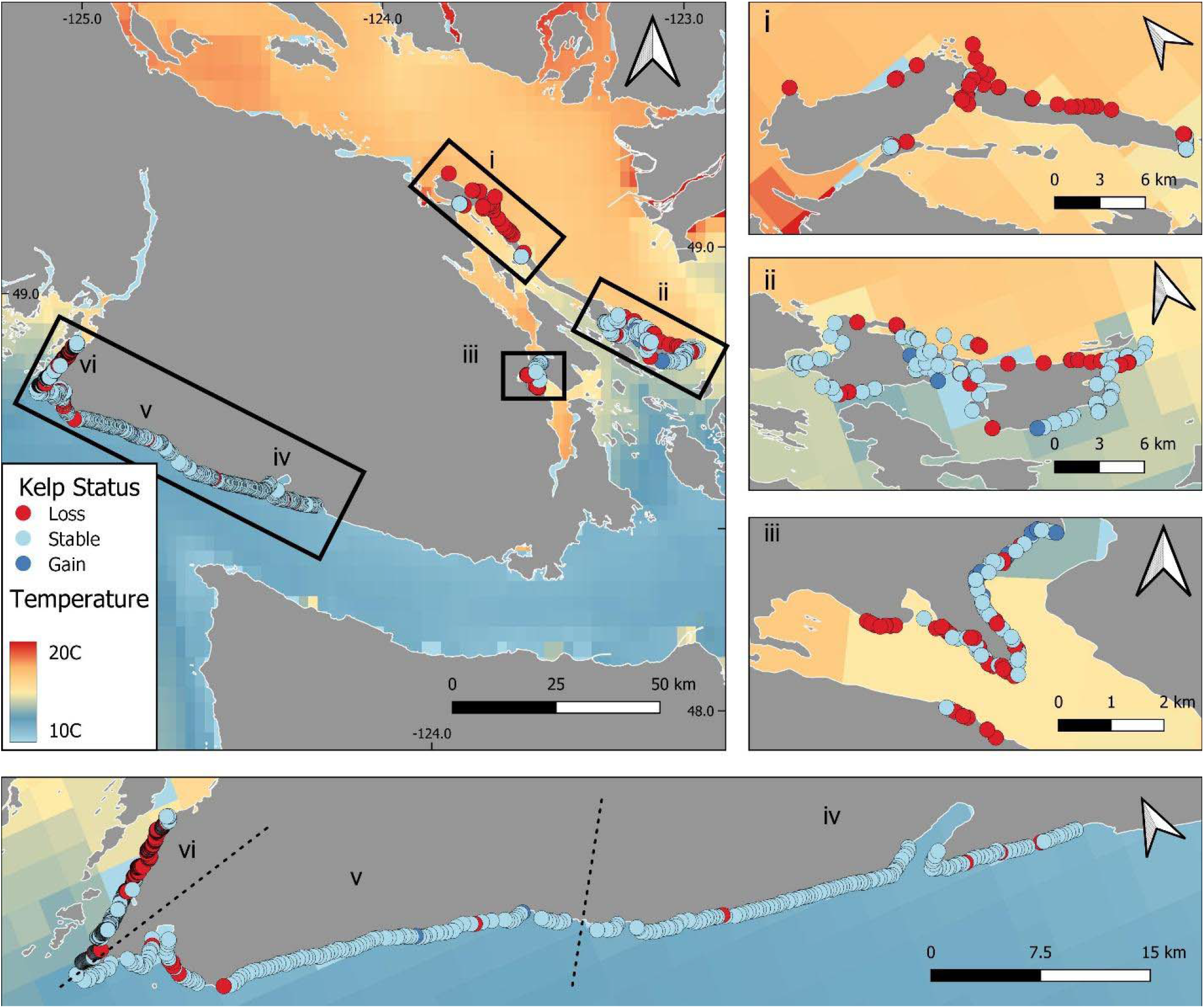
Kelp persistence within and across regions on southern Vancouver Island related to summer sea surface temperature. Data points indicate segments from the snapshot analysis coded by their persistence status (see inset legend) determined by comparing data from before (2004 – 2007) and after (2017 – 2021) the 2014-2016 MHW – see Table 1. Colored layer indicates mean sea surface temperature in August 2017, as inferred from the LiveOcean Model. Shown are both Southeast and Southwest regions of Vancouver Island (i: Valdes/Gabriola, ii: Mayne/Saturna, iii: Cowichan Bay, iv: Juan de Fuca Entrance, v: West Coast Trail, vi: Barkley Sound).

Exposure to waves and currents appeared to also influence kelp forest persistence. While there was extensive kelp loss in Valdes/Gabriola (region i), there were persistent forests in the high current narrows between islands. Similarly, in the Cowichan (iii) region, the most persistent forests were in the narrows between Vancouver Island and Saltspring Island. This likely reflects either fine-scale mixing that brings cooler waters to the surface not captured by the LiveOcean Model or else it reflects impacts of water motion through other means (e.g., impacts on urchin-kelp dynamics, growth rate). In the Laredo Sound (x) and Dundas Island (xi) regions, kelp forests largely disappeared from more sheltered areas, specifically the eastern (leeward) side of Dundas Island and Aristazabal Island, respectively (Fig 4). In both cases, urchin barrens (and in some cases, groups of sea urchins) were visible as shallow as the intertidal zone in the oblique aerial imagery (Fig S3-S6). However, kelp persisted on the side of each island that faces towards the open ocean, where wave exposure is likely to be much greater. Thus, wave action may limit the ability of urchin grazers to reach the shallowest edge of the depth range of kelp forests, thereby facilitating persistence of kelp in the face of increasing urchin populations (Keats, 1991; Watson & Estes, 2011).

**Fig 4.**
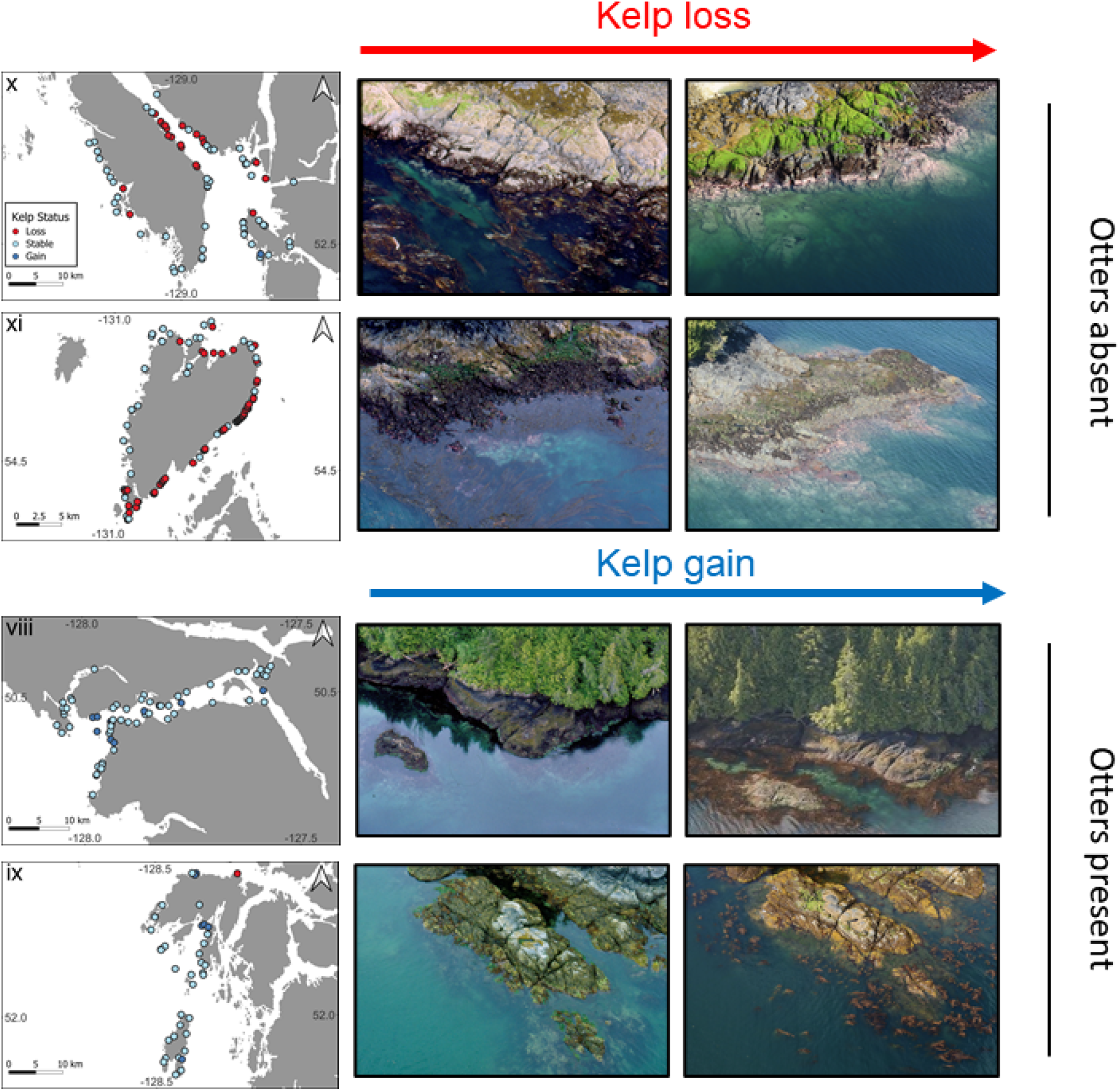
Local-scale (within-region) variation in kelp persistence between historical and modern imagery in northern regions with and without growing sea otter populations. Each data point represents a shoreline segment visible from images at both time points and colour indicates whether a shoreline segment had persistent kelp forests, forests that disappeared (loss) or colonisations of shoreline segments (gains). Aerial images show examples of kelp loss (top) or gains observed in each of these regions. Images from ShoreZone BC and Environment and Climate Change Canada.

Kelp forest gains also tended to be spatially clustered, primarily in regions with increasing sea otter populations (Fig 4). Gains in Nootka Sound were focused in two parts of the region, the inner islands of the sound and a small islet towards the exposed outer coast. Gains in Quatsino Sound tended to be spatially clustered around the opening of the sound but were also found in a few other cases further into the sound. On the South Central Coast, two stretches of coastline that included multiple shoreline segments each were colonised by kelp while remaining largely stable elsewhere. Kelp forest colonisations in other regions were generally patchy and uncommon but tended to be found nearby other segments that had kelp in both time points.

Time series from a variety of regions provide temporal context for observed changes in the distribution of kelp forests from the snapshot analyses. Across several regions with time series, negative impacts from the 2014-2016 MHW appear prevalent with varying levels of recovery and in some cases continued declines. The Central Strait of Georgia (region ii-b; only *Nereocystis* present) and Barkley Sound (region vi; both species present but *Macrocystis* more common) both experienced declines during this time-period and these have persisted for several years past the event (Fig 5a, f). In contrast, while Mayne Island (region ii) and Cowichan Bay (region iii) both had the lowest linear extent in the year following the MHW (2017) than in any other year (including during the MHW), these declines were relatively moderate (~22 and ~34% compared to lowest pre-MHW year) and for Mayne Island, recovery of the kelp forests was captured in later years. However, despite this apparent recovery in the regional Mayne Island time series, some individual sites experienced persistent declines during and following the MHW (Fig S7). Data from part of Laredo Sound (region x) demonstrate that kelp was present throughout the entire region as recently as 2013. Thus, kelp loss in Laredo Sound captured in our snapshot analysis occurred sometime between 2013 and 2019, which coincides with the timing of the MHW and SSWD impacts. Similar patterns were observed on Calvert Island (ix-b), where kelp area declined between 2015 and 2020. Interestingly, 2014 had the greatest bull kelp abundance at the North Beach site on Calvert Island but steadily decreased over the next several years. This spike in kelp abundance in 2014 coincides with short-term occupation by sea otters which occurred in this region between 2013 and 2014 (Burt et al., 2018). Importantly, kelp abundance following the departure of sea otter populations and the 2014-2016 event was lower than kelp observations beforehand (2006, 2012). Temporal variation in bull kelp abundance at this site therefore lends insight into multiple ecosystem states: kelp forests with *Pycnopodia* but no otters (2006 – 2013), kelp forests with both sea otters and *Pycnopodia* (2014), and kelp forests without either top predator (2016 – 2021) (Burt et al., 2018).

**Fig 5.**
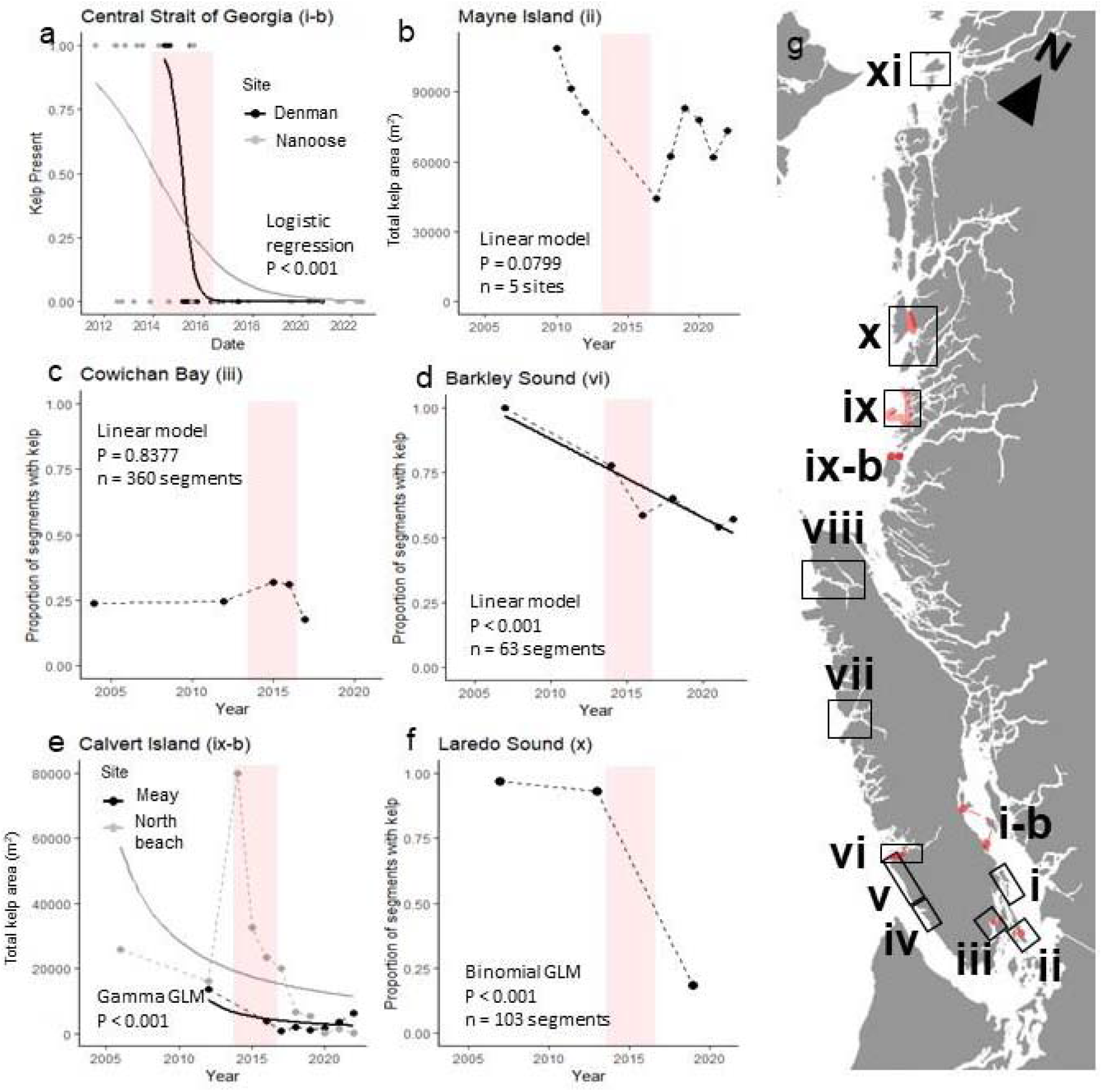
Time series from six regions spanning the British Columbia coastline. Inset map shows sites or regions used for time series data highlighted in red, relative to the snapshot regions shown with boxes. Data points indicate measures of kelp abundance or occupancy years in the time series. Note that while data points are connected by dotted lines, this does not imply continuous sampling as there are gaps in the years included in each time series (see Tables 2, S3).

The longest running time-series is that of total offshore kelp area for the South Central Coast (ix). This is a region that was colonised by sea otters around the year 1990 and otter populations have been increasing since (Nichol et al., 2015, 2020). Time series data from this region do not include kelp within 30m of shore but offer an overall look at the abundance of kelp in offshore beds (e.g. >30m from shore) which are common in this area. The time series shows no directional trend through time (Linear model: P = 0.99) with some evidence of increase in kelp abundance early in the dataset. Specifically, the highest abundances were found between 1999 and 2014 despite the dataset dating to 1984. Although kelp abundance was approximately average in 2014, it dropped to its lowest value in 2015. While 2015 kelp area was only <5 % less than multiple other years of the dataset, recovery following the 2014-2016 MHW was limited. Kelp abundance in 2015 and later tended to be lower than the mean kelp abundance in years prior, however this trend was not significant (W = 158, p = 0.06501). While past declines in kelp abundance to similar levels were generally followed by rapid recovery, hinting at resilience in these forests, kelp abundance following the event has stayed reduced for another six years following the initial crash in 2015. It is worth noting that declines in kelp area captured in the time series are not reflected in the snapshot analysis. Because kelp forests in the snapshot analysis are generally fringing beds within 30m of the shoreline that are excluded from the satellite time-series, this may reflect either differences in the type of kelp forest assessed by the two methods or differences in the metrics used (i.e., shoreline occupancy vs. total area). This highlights that changes in linear extent and total kelp area can be decoupled in some cases and underscores the importance of examining differences in kelp stability according to local and regional spatial scales as well as bed morphology.

## 4. Discussion

Here, we established for the first time that kelp forests in British Columbia have experienced variable patterns of change over the timeframe of recent marine heatwave events, with many regions experiencing substantial declines in kelp linear extent, consistent with our first hypothesis. In the most extreme cases, kelp extent declined by more than 60% in two regions (Valdes/Gabriola (i): 74% loss; Dundas Island (xi): 62% loss) and more than 30% in two others (Barkley (vi): 43% loss; Laredo Sound (x): 31% loss) in the snapshot analyses, supporting our second hypothesis of near-complete loss in some regions. While our snapshot analyses were not comprehensive in coverage of coastal British Columbia, kelp declines were observed across the coastline; with modest increases (14 – 22 % gains) observed in only two regions and little to no change observed in three regions, hinting at regional-scale kelp refugia. Moreover, where snapshot and time series data overlap, declines in kelp abundance and biomass are unprecedented over the time periods of time series. For example, even the highly variable time series of the South Central Coast (region ix) shows reduced resilience (i.e., ability to recover following a year with low kelp abundance) following the 2014-2016 marine heatwave (MHW) compared to the three decades prior. Similarly, kelp reached its lowest occupancy states in Barkley (vi) and Laredo (x) Sounds following the heatwave compared to past timepoints spanning multiple decades (Fig. 5; see Starko et al. 2022 for in depth case study of Barkley Sound to the 1970s). Overall, the scale and persistent nature of concurrent kelp forest loss across several B.C. regions suggests that kelp forests should be a conservation concern in this province. Importantly, however, not all regions have experienced losses and instead declines have been spatially clustered to particular localities within only some regions. This highlights how ecosystem-level perturbations have different impacts depending on the underlying heterogeneity in the environment, a phenomenon that should be strongly considered when developing management plans and monitoring programs.

### 4.1 Spatial variation in kelp forest trajectories

Patterns of change were strongly spatially structured, providing insight into the drivers behind those changes. In areas that are naturally warmer due to seasonal patterns of warming (e.g., Central Strait of Georgia (i-b), Gabriola/Valdes (i), inner parts of Barkley Sound (vi)), kelp forests largely disappeared during the 2014-2016 MHW with very minimal (if any) recovery. In contrast, nearby regions (e.g., Mayne/Saturna (ii) and the West Coast Trail (v)) experienced little net kelp loss, instead maintaining extensive kelp forests (Fig 4). These patterns are consistent with our third hypothesis and are captured in both snapshot imagery analysis and time series data, where the timing of kelp declines in both Barkley Sound and the Central Strait of Georgia coincided with the 2014-2016 MHW (Fig 5). Although it is challenging to disentangle the direct impacts of temperature from those of urchin expansions expected from the die-off of *Pycnopodia*, recent work in Barkley Sound demonstrated that these factors together can drive kelp loss in warm areas by negatively impacting kelp forests across their depth range, preventing persistence in both shallow and deeper waters (Starko et al., 2022).

We attribute the loss of kelp in the two northern-most regions without otters (Laredo Sound – region x and Dundas Island – region xi) to increases in urchin grazing. Both of these regions experienced large declines in kelp extent, despite no evidence that these regions experience temperatures warm enough to threaten kelp forests. Lighthouses nearby to both regions show that temperatures remained consistently below 18°C during the summers of 2014-2017 (Fig 1) which include the warmest years in decades (see Fig 5). Instead, oblique imagery clearly shows that kelp forests in these regions have transitioned to urchin barrens, with sea urchins visible as shallow as the intertidal zone along several stretches of coastline where kelp has disappeared (Figs 3, S4). Kelp losses in these areas are largely restricted to the leeward side of islands, where the coast is sheltered from incoming swell. In contrast, coastlines facing west in both regions generally retained kelp forests. Wave action from oncoming swell can limit the depth that sea urchins can graze (Keats, 1991; Kawamata, 2010; Watson & Estes, 2011), potentially facilitating kelp forest persistence in shallow waters, despite increases in the abundance and dominance of sea urchins. Indeed, the West Coast Trail (v) and Juan de Fuca (iv) regions, which both face the dominant direction of oncoming swell and are generally cooler due to mixing at the entrance of the Juan de Fuca Strait (Fig 1; see station 1), experienced very little (<5%) change in kelp extent, likely due to the absence of both environmental and biotic drivers of decline. Past work has suggested that tidally driven vertical mixing might allow these regions to serve as climatic refugia for marine systems in the face of climate warming and perturbations (Ban et al., 2016). This hypothesis is strongly supported by our results.

The only regions to experience more increases in kelp extent than decreases were those with growing sea otter populations (regions vii to ix), consistent with our fourth hypothesis that these regions would be more resistant and/or resilient to changes in trophic dynamics. Although sea otters were once widely distributed in British Columbia, they were extirpated from the entire coast during the Fur Trade of the 18^th^ to 20^th^ centuries (Nichol et al. 2015). Sea otters were reintroduced to Checleset Bay on Northern Vancouver Island in 1969 - 1972 and have since expanded to include all three of regions vii to ix included in our study. Moreover, provincial sea otter surveys conducted as recently as 2017 indicate that otter population sizes have continued to increase in all study regions in which they have re-established (Nichol et al. 2020). Thus, increases in kelp extent in these regions likely reflect successional dynamics associated with changes in trophic structure (Watson and Estes 2011). Specifically, increasing otter populations would be expected to drive declines in sea urchin abundance which could subsequently allow kelp to colonise stretches of shoreline that were previously in the urchin barren state. We note that patterns of persistence and colonisation are similar across both canopy-forming kelp species in regions where the two species co-occur (Fig. S8).

### 4.2 Timing of kelp forest change

Time series data from seven regions capture widespread declines between 2014 and 2017, coincident with the MHW and SSWD event. The Central Strait of Georgia (i-b), Barkley Sound (vi), Calvert Island (ix-b) and Laredo Sound (x) all show evidence of persistent declines, with little to no recovery following the heatwave (Fig 5). Where data span multiple years of the heatwave (regions i-b, iii, vi, ix-b), losses were sometimes not documented until the second year of the MHW or later (2015-2017), suggesting that the multi-year nature of the event was critical to driving declines. In contrast to regions with persistent declines, kelp forests around Mayne Island (region ii) experienced negative impacts from the 2014-2016 marine heatwave but these declines were not persistent and kelp forests largely recovered in following years. However, site-level analyses around Mayne Island (Fig S7) indicate that the trajectories of individual kelp forests have been variable with some sites remaining stable or increasing and others experiencing persistent declines. This highlights how variation in kelp forest trajectories often occurs at fine scales with apparent site-level differences in the persistence of kelp forests.

### 4.3 Caveats and future directions

Kelp forests are naturally highly variable systems that tend to fluctuate interannually, a pattern clearly demonstrated by our analysis of kelp abundance in the South Central Coast (ix) time series. For this reason, there are potentially important caveats associated with our snapshot analyses of two time points. In particular, the timing of both historical and modern imagery has the potential to produce misleading results under some circumstances. For example, historical snapshot data from the South Central Coast (ix) region were from 1997 which was during a large marine heatwave (1997-1998 El Nino), an event which was known to negatively impact kelp in California (e.g., Ladah & Zertuche-González, 2004; Edwards & Hernandez-Carmona, 2005) and apparently drove declines in offshore kelp abundance on the South Central Coast of B.C. (Fig 6). Thus, the timing of this initial survey has the potential to bias patterns towards perceived increases. Importantly, however, pre-MHW imagery from nearby Laredo Sound (x), a region with evidence of strong declines and no sea otter populations, was also taken during the 1997-1998 event (Table 1, Fig S9). This suggests that these two regions have, in fact, experienced differing trajectories and that historical sampling during the 1997-1998 event does not necessitate a perception of kelp extent increases.

**Fig 6.**
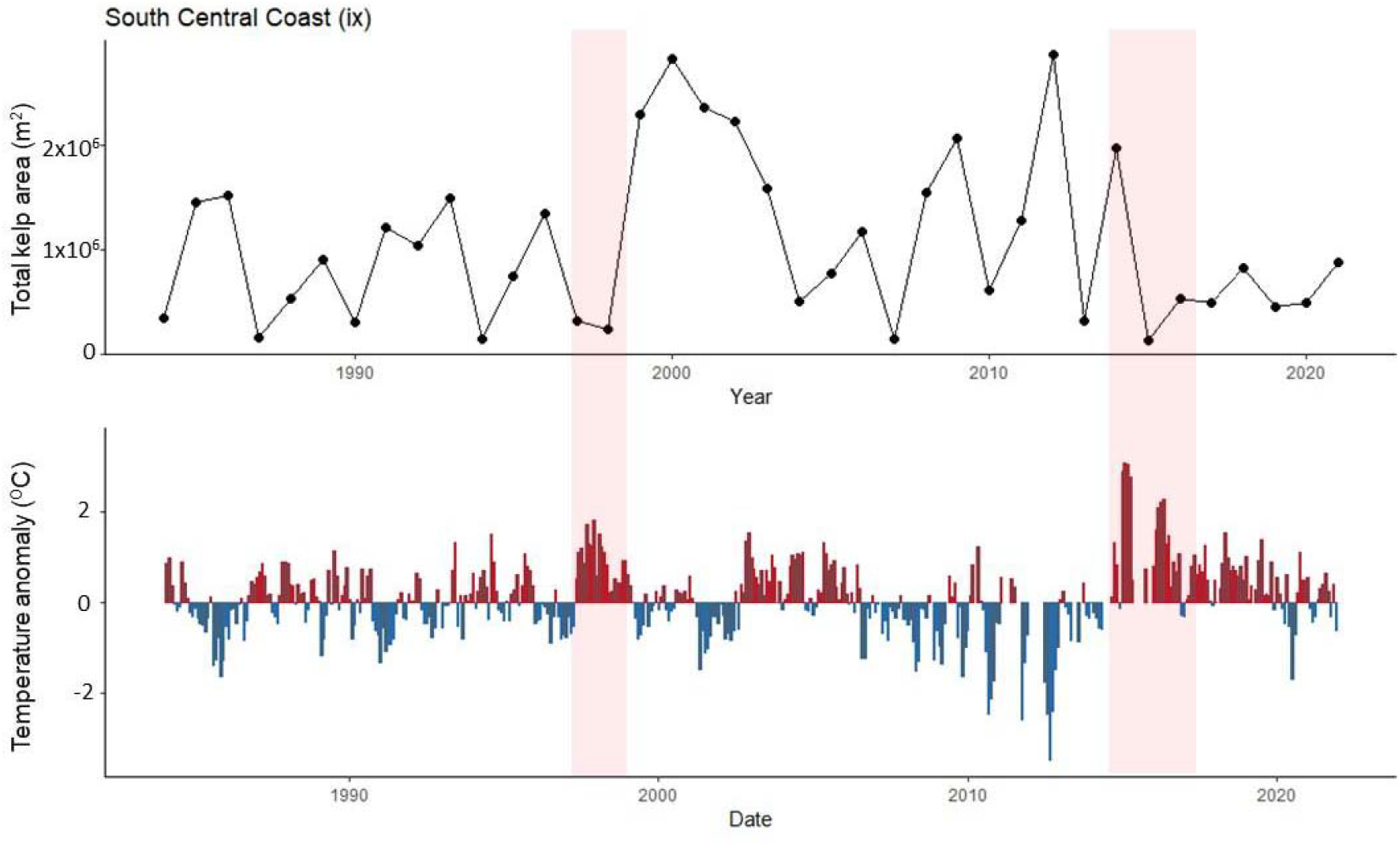
Time series of offshore kelp forest area from South Central Coast region showing high resilience prior to the 2014-2016 heatwave. (a) Data were generated from LandSat imagery and exclude kelp forests within 30m of the shore. (b) Temperature measurements from nearby McInnes Island lighthouse. Red shading highlights the timing of the 1997-1998 El Niño and 2014-2016 marine heatwave.

Similarly, due to slight differences in the timing of imagery, seasonal patterns (e.g., timing of annual canopy reaching the surface or increasing canopy biomass during the growing season) may confound true interannual patterns of change. Quatsino Sound (viii) was sampled initially in May but resampled in July when kelp beds are expected to be larger, creating potential for seasonal patterns to confound true patterns of change in kelp extent (more kelp expected later in the summer). However, the South Central Coast (ix) region had the opposite issue (with initial surveys occurring later in the season than modern imagery) and showed similar patterns of extent increases as observed in Quatsino. Moreover, all other regions had very closely matched dates for historical and modern imagery (i.e., less than one month and generally within 1-2 weeks).

Interannual variation in kelp abundance driven by environmental conditions, for example associated with climatic oscillators (e.g., PDO, ENSO), was also not captured using our two time point snapshot analyses, introducing an additional caveat to the interpretation of these data. Five of the regions examined in the snapshot analyses were also considered using time series, as well as region i-b which is adjacent to a sixth snapshot region (i.e., region i). These time series support the hypothesis that declines occurred during the 2014-2016 MHW and provide additional temporal context to patterns observed in a majority of regions. Moreover, snapshot comparisons in the Strait of Georgia (regions i, ii, iii) were all made between 2004 and 2017-2021. In 2004, PDO, ENSO and temperature anomalies were all positive (at times more so than resurvey years; Fig S9), suggesting that these two time points should experience similar effects of climatic oscillators. Yet, patterns of kelp loss at the latter sampling point were dramatic (e.g., Valdes/Gabriola lost kelp in 74% of segments). Snapshot analyses also focused only on presence-absence and not on abundance which is generally expected to be more stable even in ecologically dynamic systems (Wilson 2012). Moreover, most snapshot data were derived from oblique imagery which was generally high enough resolution to detect even a single kelp individual, further limiting the importance of interannual variability in abundance in our presence-absence analyses. In a recent paper focused on Barkley Sound (region vi) (Starko et al., 2022), presence-absence in these shoreline units was generally consistent in years prior to the MHW, even when comparing to data from the 1970s and 1980s, despite likely interannual variation in the abundance (i.e., total area) of kelp within these beds. Similarly, for Laredo Sound, all three time points (1997, 2007, 2013) prior to the heatwave show consistent presence-absence across most shoreline units, despite variability in interannual conditions across these three years. In contrast, however, data from Cowichan Bay show much more interannual variability in kelp presence-absence. Thus, the background variation in kelp presence-absence likely varies across regions.

The dates of both pre- and post-MHW snapshot imagery vary across regions, yet the data tell a consistent story in line with our hypotheses; declines in most areas without otters (especially areas of warming) and stability or increases in places with sea otters or persistent mixing. Thus, in the regions where widespread declines occurred, they were likely not restricted to a single year despite our use of only two time points to characterise them. Overall, these factors make us confident that declines observed in our snapshot analysis describe true changes in kelp linear extent with important ecological and conservation implications.

For both snapshot analyses and time series data, the data type used may have also influenced the observed patterns. Where shoreline segments were used, variation in their length may have influenced the patterns inferred from these analyses. In particular, absence observations are more likely when segments are shorter. Importantly, regions were mostly compared only to themselves, and segment length was consistent through time within each region. Moreover, in Laredo Sound, where two different segment methods were used (variable length for snapshot analysis, fixed length of 50m for time series), the same pattern was recovered in both cases, suggesting that these minor differences in length did not impact inferred patterns. The Cowichan Bay time series had larger segments (100m), which would make absences less likely. However, counter to expectation, this region was the most variable through time in terms of presence-absence along segments. Thus, this variability cannot be explained by segment length. It is worth noting that because Cowichan Bay data were derived from high resolution satellite rather than oblique aerial imagery, small fringing beds may have been classified as kelp absence points (see discussion of accuracy in Schroeder et al. 2019). Further, this region is characterized by high currents which can easily submerge fringing kelp and reduce the ability to detect it at the surface (Britton-Simmons, Eckman & Duggins, 2008; Timmer et al., 2022). Thus, false negatives are probably more likely in this one region than in other regions analyzed using two time points. For the time series analyses, a number of different survey methods and response variables were assessed based on the availability of data. While we may expect these different metrics to be sensitive to different types of patterns, it is important to note that time series were not compared to each other but rather used to assess changes through time within a region.

Future work should aim to expand on this study in multiple ways. Firstly, the growing availability of satellite imagery (including some products dating back decades) will allow researchers to reconstruct time series in regions for which they are not already available (Cavanaugh et al., 2021; Gendall, 2022), especially in regions with large offshore beds (Nijland et al. 2019). This approach may help facilitate a province-wide assessment of canopy kelp persistence, rather than focusing on a subset of regions as we have done here. Alternatively, qualitative approaches may also be useful in assessing the extent of kelp forest loss in some regions. For example, Traditional and Local Ecological Knowledge could help to identify regions of major change for which there is no alternative data or could supplement and increase confidence in quantitative approaches (Reid et al., 2021). Similarly, herbarium records (Wernberg et al., 2011) and historical nautical charts (Costa et al., 2020) may offer insights into the historical distribution of kelp species, especially if records are available from regions that no longer support any kelp forests. Finally, in the face of environmental change, it will be essential to not only reconstruct past kelp forest distributions but also make predictions about future change. This can be accomplished by coupling species distribution models with climate projections that could help to identify areas of resilience or vulnerability in the face of global change (e.g., Martínez et al., 2018; Chefaoui et al., 2021b).

### 4.4 Conclusions and implications

Here we showed that kelp forests have experienced variable patterns of change across coastal British Columbia, with recent and substantial declines in some focal regions. Declines in kelp forest linear extent and/or abundance appear linked to both rapid warming experienced during and after the prolonged 2014-2016 MHW and to increases in herbivorous urchins driven by the loss of *Pycnopodia* sea stars. Importantly, microclimate, wave exposure and food chain length strongly mediated the impacts of these drivers on kelp forest ecosystems in B.C., causing some regions to be particularly sensitive to these drivers while others remained stable or increased, indicative of climatic refugia. Large-scale concurrent evidence of declines suggests that kelp forest ecosystems in B.C. should be of significant conservation concern across much of the province. However, to be effective, conservation and management efforts should focus on parts of the coast that are most sensitive to environmental and biological drivers of change, rather than treating kelp forests across all regions as equally sensitive to environmental change. Overall, our findings highlight how local or regional scale conditions can be essential in determining the impacts of extreme warming events on coastal marine ecosystems and demonstrate that kelp forest loss in B.C. offers a major conservation challenge in the face of ongoing global change.

## 5. Acknowledgements

The authors extend our deep gratitude to the numerous First Nations in whose traditional, unceded or treaty lands and waters this work was conducted. We are grateful for our existing partnerships with First Nations groups at the time of manuscript preparation and hope that it will set the stage for additional collaborations with other Nations with interest in furthering our collective understanding of these ecosystems that they have long stewarded and on which we all rely. S. Starko and J. Baum acknowledges funding from Mitacs; S. Starko and C. Neufeld acknowledge funding from the Ngan-Page Family Fund; S. Starko, M. Costa, S. Schroeder, W. Heath, A. Zielinski, and R. Zielinski acknowledge funding from the Pacific Salmon Foundation; J. Baum and M. Costa acknowledge funding from the Natural Sciences and Engineering Research Council. M. Hessing-Lewis and L. Reshitnyk thank the Tula Foundation for funding long term ecological research and monitoring conducted by the Hakai Institute on the Central Coast. W. Heath, A. Zielinski and R. Zielinski acknowledge the Fisheries and Oceans Canada Coastal Restoration Fund. We thank the ShoreZone initiative and Environment and Climate Change Canada for providing access to imagery used for a substantial proportion of our analyses. We thank Parks Canada and the Huu-ay-aht First Nations for contributing to the funding of some ShoreZone flights in 2021. We thank Mark Bright for contributing Nanoose SCUBA videos used in the analyses. We thank support from the Bamfield Marine Sciences Centre, the Mayne Island Conservancy and all staff employed therein.

