## Supplementary Materials for "Temperature and food chain length, but not latitude, explain region-specific kelp forest responses to an unprecedented heatwave"

**Running Head:** Kelp forest responses to prolonged heatwave

Samuel Starko, Brian Timmer, Luba Reshitnyk, Matthew Csordas, Jennifer McHenry, Sarah Schroeder, Margot Hessing-Lewis, Maycira Costa, Amanda Zielinksi, Rob Zielinksi, Sarah Cook, Rob Underhill, Leanna Boyer, Christopher Fretwell, Jennifer Yakimishyn, William A. Heath, Christine Gruman, Julia K. Baum & Christopher J. Neufeld

Table S1. Sources of data used in the snapshot analysis. ECCC = Environment and Climate Change Canada. MP = *Macrocystis pyrifera*; NL = *Nereocystis luetkeana.*

| Region | Timepoint 1 (T1) | Data source (T1) | Timepoint 1 (T2) | Data source (T2) | Kelp species present |
| --- | --- | --- | --- | --- | --- |
| Valdes and Gabriola Islands (i) | July 29, 2004 | ShoreZone | August 7, 2021 | ShoreZone | NL |
| Mayne and Saturna Island (ii) | July 29, 2004 | ShoreZone | August 7, 2021 | ShoreZone | NL |
| Cowichan Bay (iii) | September 24, 2004 | QuickBird Satellite | July 27, 2017 | WorldView3 Satellite | NL |
| Juan de Fuca Entrance (iv) | August, 13-14, 2007 | ShoreZone | August 9, 2021 | ShoreZone | NL |
| West Coast Trail (v) | August 14, 2007 | ShoreZone | August 8-9, 2021 | ShoreZone | MP, NL |
| Barkley Sound (vi) | August 14, 2007 | ShoreZone | August, 2018 | In situ survey | MP, NL |
| Nootka Sound (vii) | June 26, 1994 | ShoreZone | July 24, 2021 | ShoreZone | MP, NL |
| Quatsino Sound (viii) | May 17, 1999 | ShoreZone | June 17, 2018 | ECCC | MP, NL |
| SouthCentral Coast (ix) | July 21, 1997 | ShoreZone | May 18, 2018 | ECCC | MP, NL |
| Laredo Sound (x) | July 24, 1997 and July 12-13, 1998 | ShoreZone | July 7, 2019 | ECCC | MP, NL |
| Dundas Island (xi) | July 2, 2000 | ShoreZone | July 4, 2019 | ShoreZone | MP, NL |

Table S2. Years and data sources for time series from the Central Strait of Georgia

| Site | Year | Number of dives | Source |
| --- | --- | --- | --- |
| Eagle Rock, Denman Island | 2014  2015  2016  2017  2020  2022 | n = 4  n = 4  n = 7  n = 2  n = 2  n = 1 | Project Watershed and Hornby Island Divers |
| Tyee Cove, Nanoose Bay | 2011  2012  2013  2014  2015  2016  2017  2019  2020  2021  2022 | n = 1  n = 5  n = 4  n = 3  n = 3  n = 4  n = 2  n = 2  n = 3  n = 3  n = 4 | Mark Bright, Dean Driver, SCUBA BC, Rowan Costall, Rayce Bannon, Tom Hlavac, Gerald Huppertz, Mike Holmes, Oceanside, Mark Heibert |

Table S3. Sources of data used in time series analyses for all regions except the Central Strait of Georgia.

| Region | Year | Data type | Source | Methods reference (if applicable) |
| --- | --- | --- | --- | --- |
| Mayne Island (iii) | 2010, 2011, 2012, 2017, 2018, 2019, 2020, 2021 | *In situ* | Kayak surveys (Mayne Island Conservancy) |  |
| Cowichan Bay (iii) | 2004 | Satellite | QuickBird | Schroeder et al. 2020 |
|  | 2012 | Satellite | WorldView 2 | Schroeder et al. 2020 |
|  | 2015 | Satellite | WorldView 2 | Schroeder et al. 2020 |
|  | 2016 | Satellite | WorldView3 | Schroeder et al. 2020 |
|  | 2017 | Satellite | WorldView3 | Schroeder et al. 2020 |
| Barkley Sound (vi) | 2007 | Oblique aerial imagery | Shorezone | Starko et al. 2022 |
|  | 2013 | Satellite | Google Earth | Starko et al. 2022 |
|  | 2014 | Aerial imagery | Air phot from GeoBC | Starko et al. 2022 |
|  | 2016 | Satellite | Google Earth | Starko et al. 2022 |
|  | 2018 | *In situ* | Boat surveys | Starko et al. 2022 |
|  | 2021 | *In situ* | Boat surveys | Starko et al. 2022 |
|  | 2022 | *In situ* | Boat surveys |  |
| South Central Coast (ix) | 1984 -2021 (continuous) | Satellite | LandSat (Google Earth Engine Kelp Tool) | Nijland et al. 2019 |
| Calvert Island (ix- b) | 2006 -2021 | Aerial imagery | RPAS |  |
| North Beach | 2006, 2012, 2014, 2015, 2016, 2017, 2018, 2019, 2020, 2021. 2022 |  |  |  |
| Maey Channel | 2012, 2016, 2017, 2018, 2019, 2020, 2021, 2022 |  |  |  |
| Laredo Sound (x) | 2007 | Aerial imagery | Air photo from GeoBC |  |
|  | 2013 | Satellite | Google Earth |  |
|  | 2019 | Oblique aerial imagery | Environment and Climate Change Canada |  |

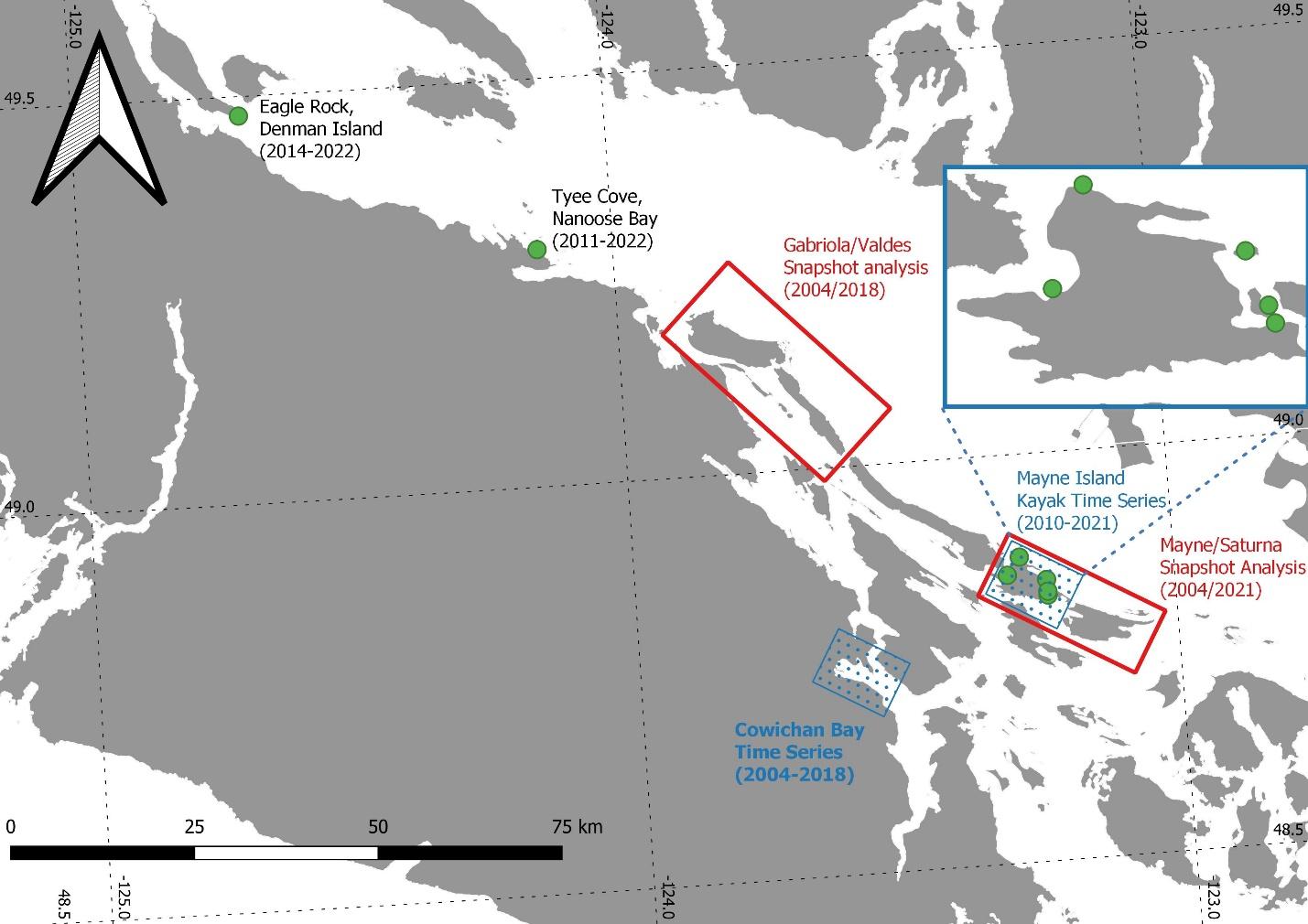

Fig S1. Map showing the locations of time series regions (blue/green) relative to snapshot analysis regions (red) for the Strait of Georgia.

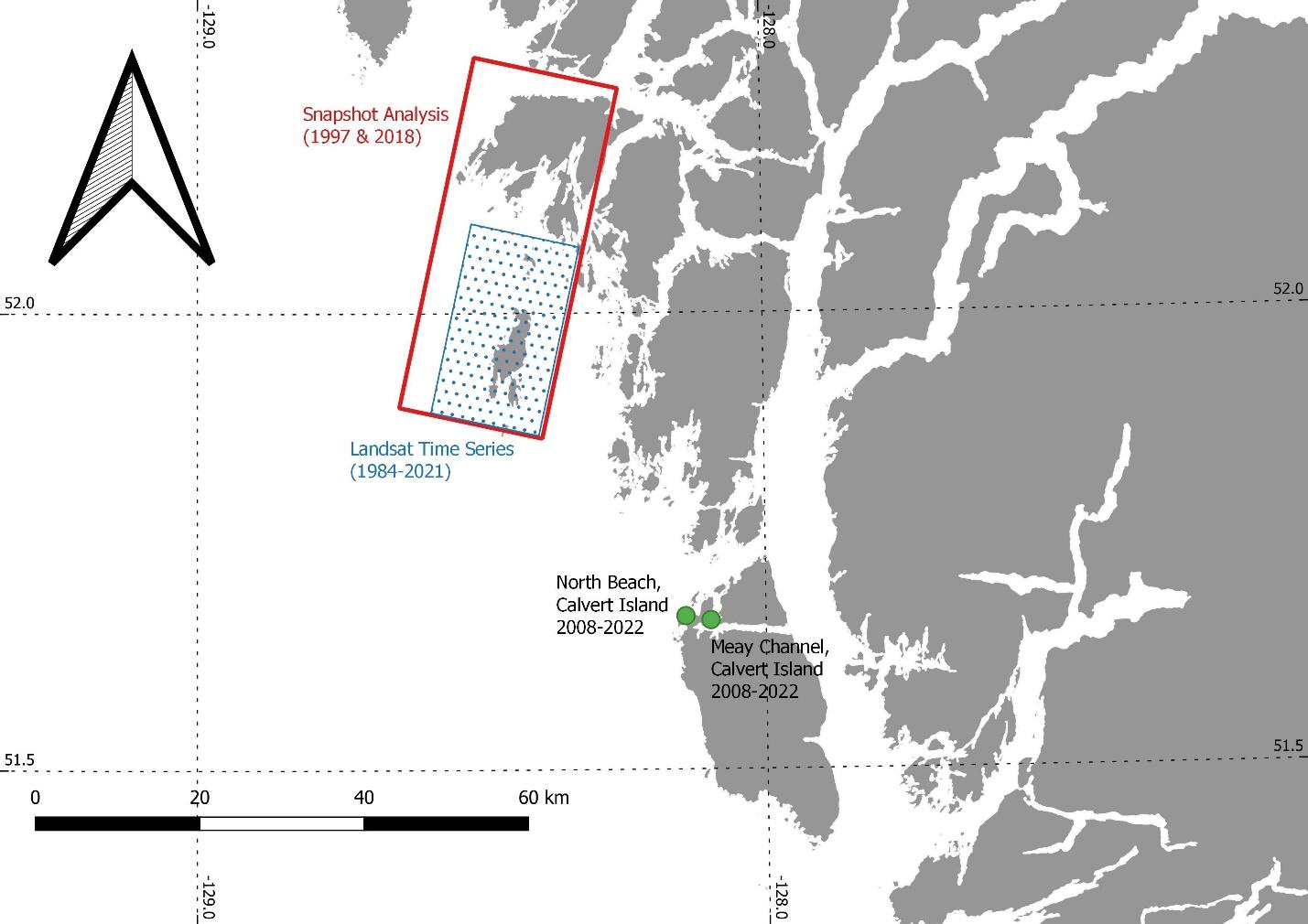

Fig S2. Map showing the locations of time series regions (blue/green) relative to snapshot analysis regions (red) for the South Central Coast and Calvert Island regions.

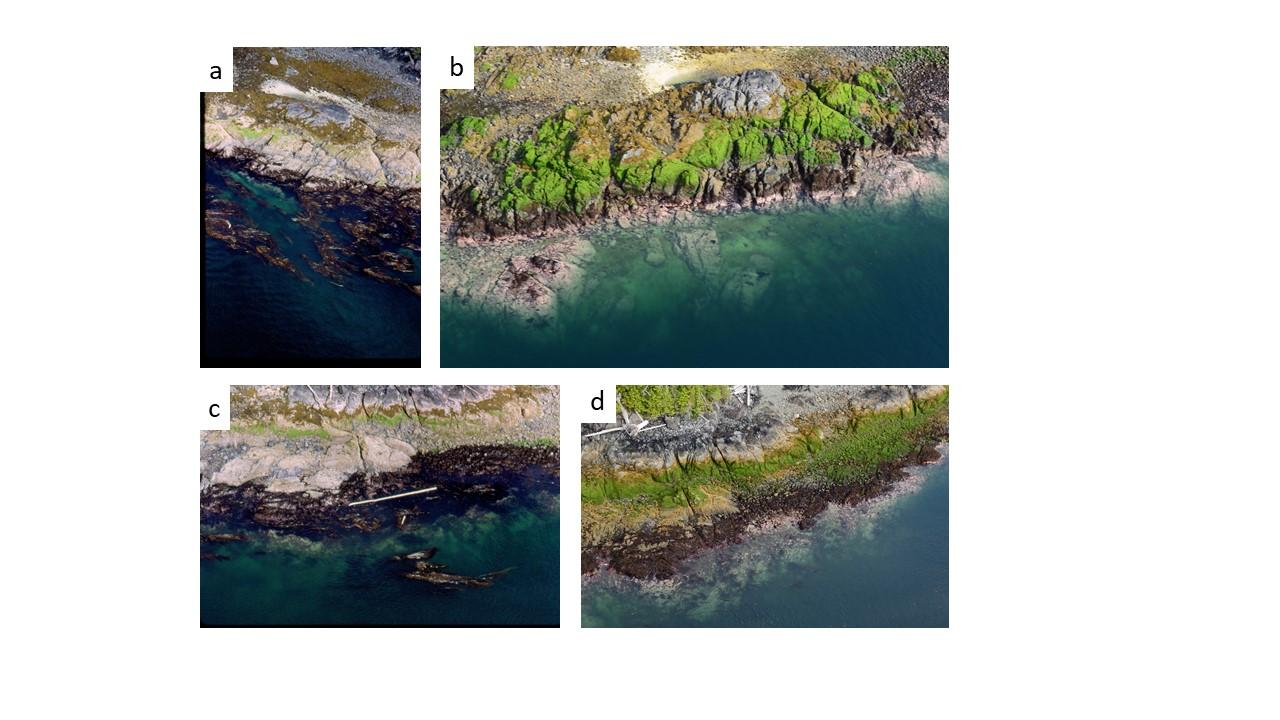

Fig S3. Examples of comparisons between modern and historic imagery from Dundas Island. Panels a and c show kelp forests in 2000 while panel b shows a transition to urchin barren and panel d shows a much reduced sparse forest surrounded by urchin barren. Images from ShoreZone BC.

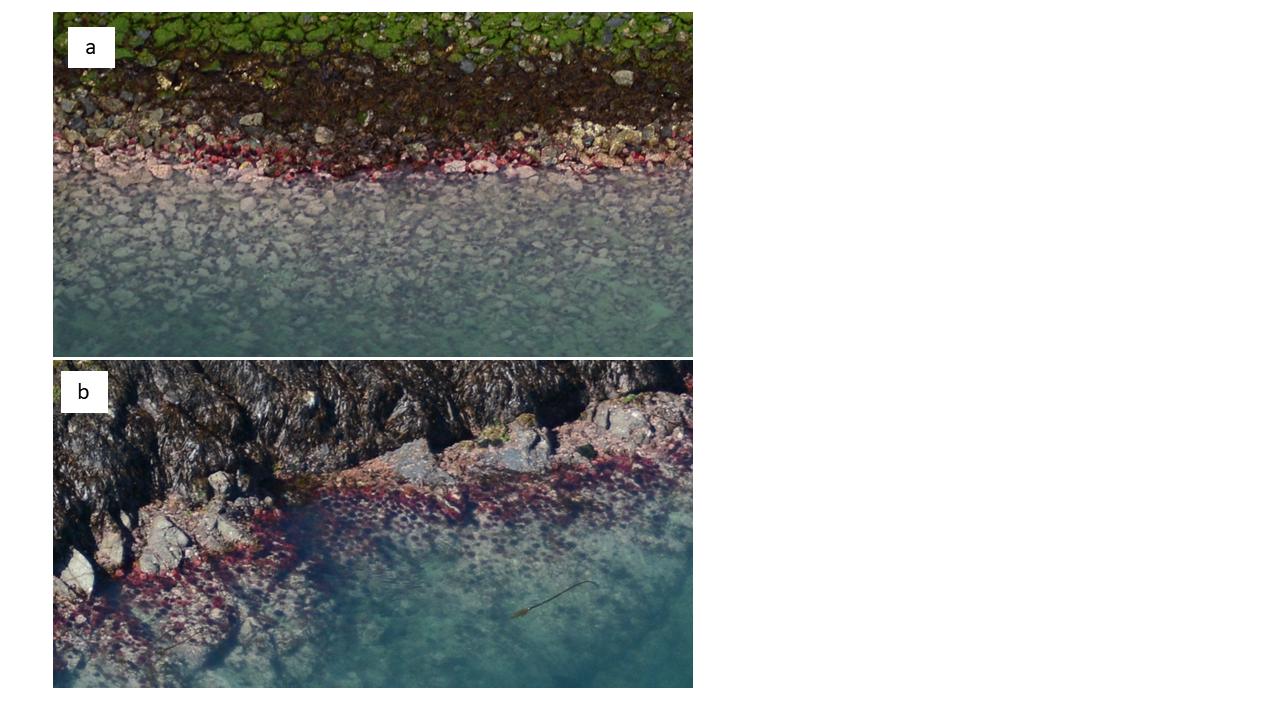

Fig S4. Example of sea urchins grazing up into the intertidal zone on Dundas Island (a) and in Laredo Sound (b) in 2019. Images from ShoreZone BC and Environment and Climate Change Canada.

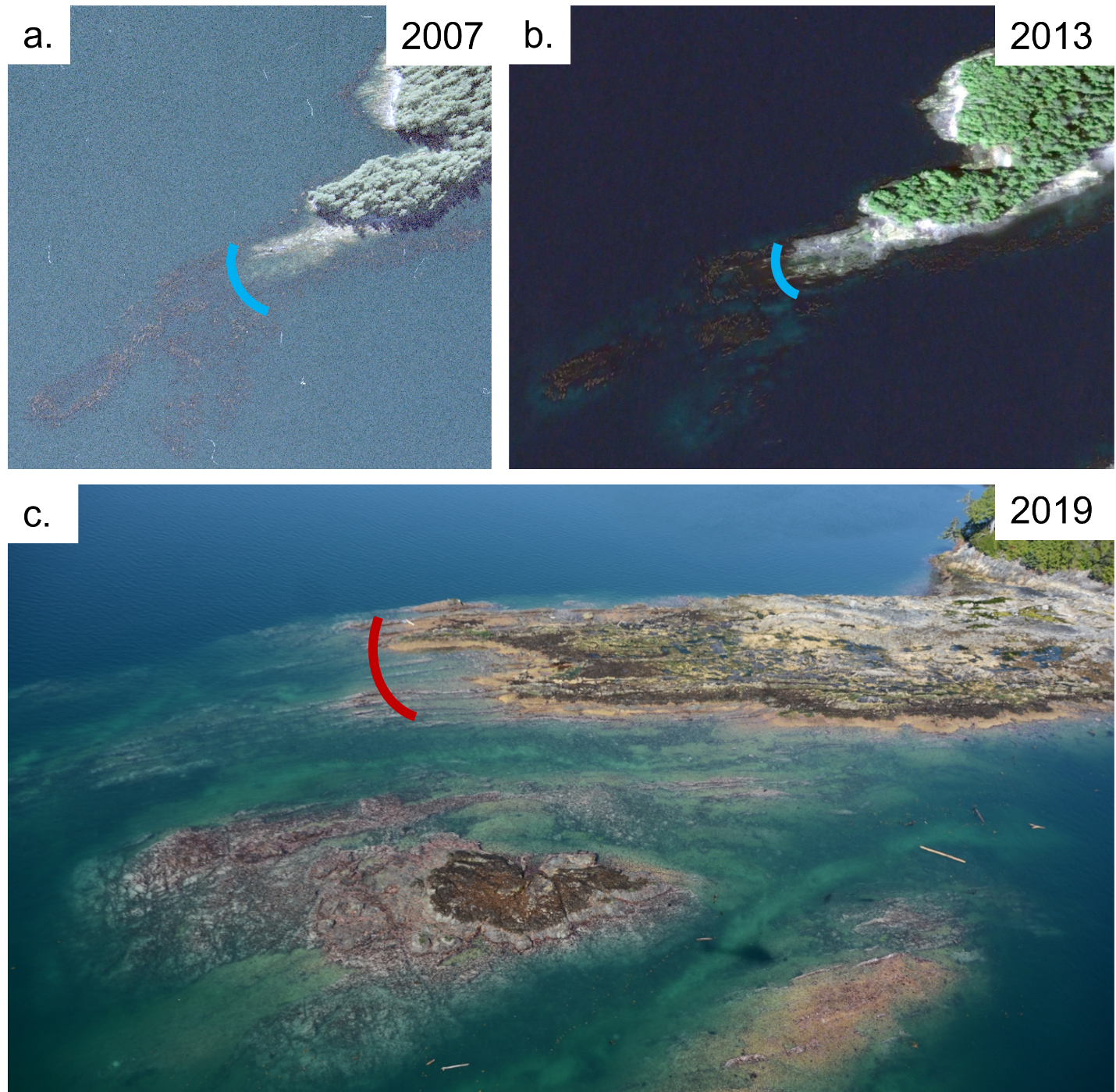

Fig S5. Examples of transitions from kelp forests to urchin barrens in Laredo Sound. Line segments show the same example area compared over time, with blue indicating kelp presence and red indicating kelp absence. Images from GeoBC, Google Earth and Environment and Climate Change Canada.

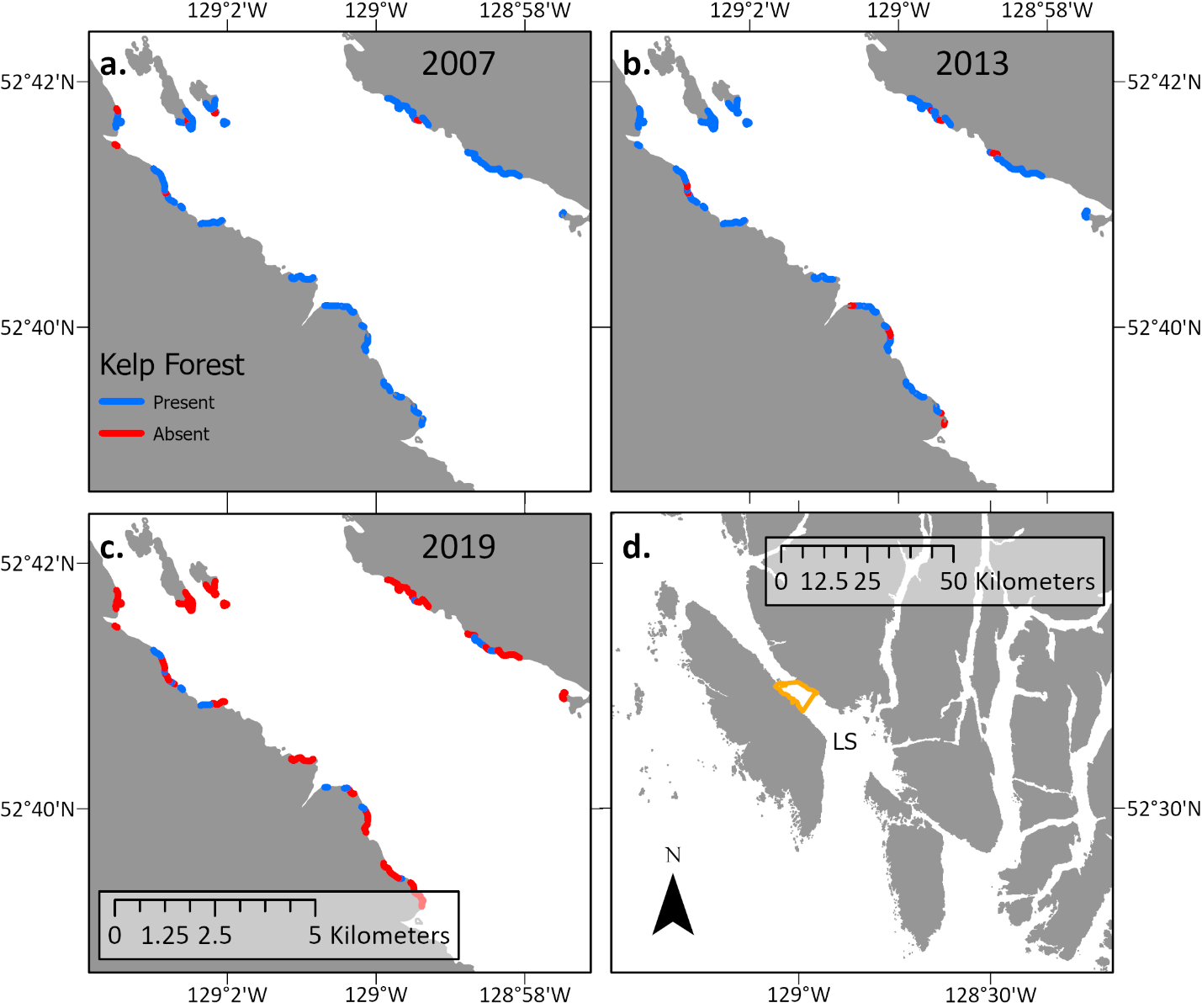

Fig S6. Maps showing the loss of kelp in a subregion of Laredo Sound between a) 2007, b) 2012, and c) 2019. Inset map shows the location of the spatial domain of the comparison within Laredo Sound.

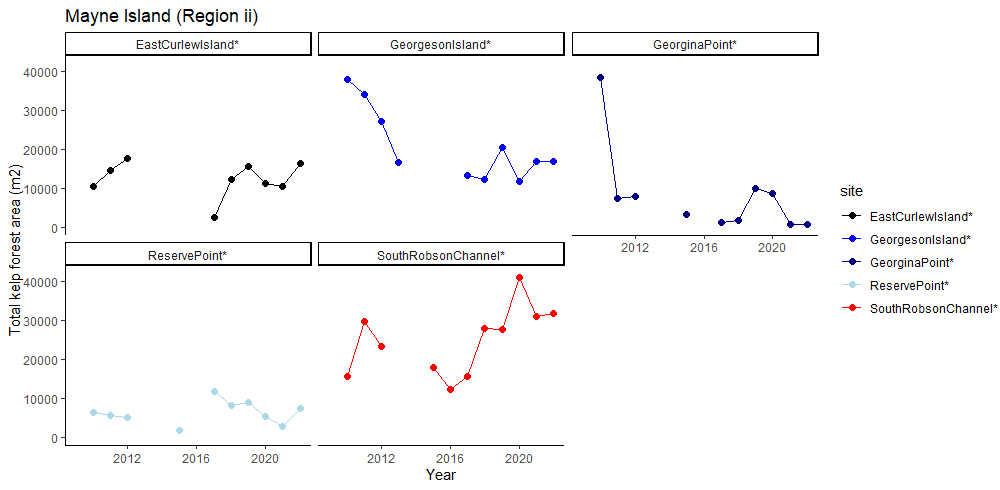

Fig S7. Site level time series of in situ monitoring data collected on Mayne Island (Region ii). Shown is kelp forest area, as inferred from in situ surveys from kayak, for each of five sites. Adjacent years are connected with solid lines.

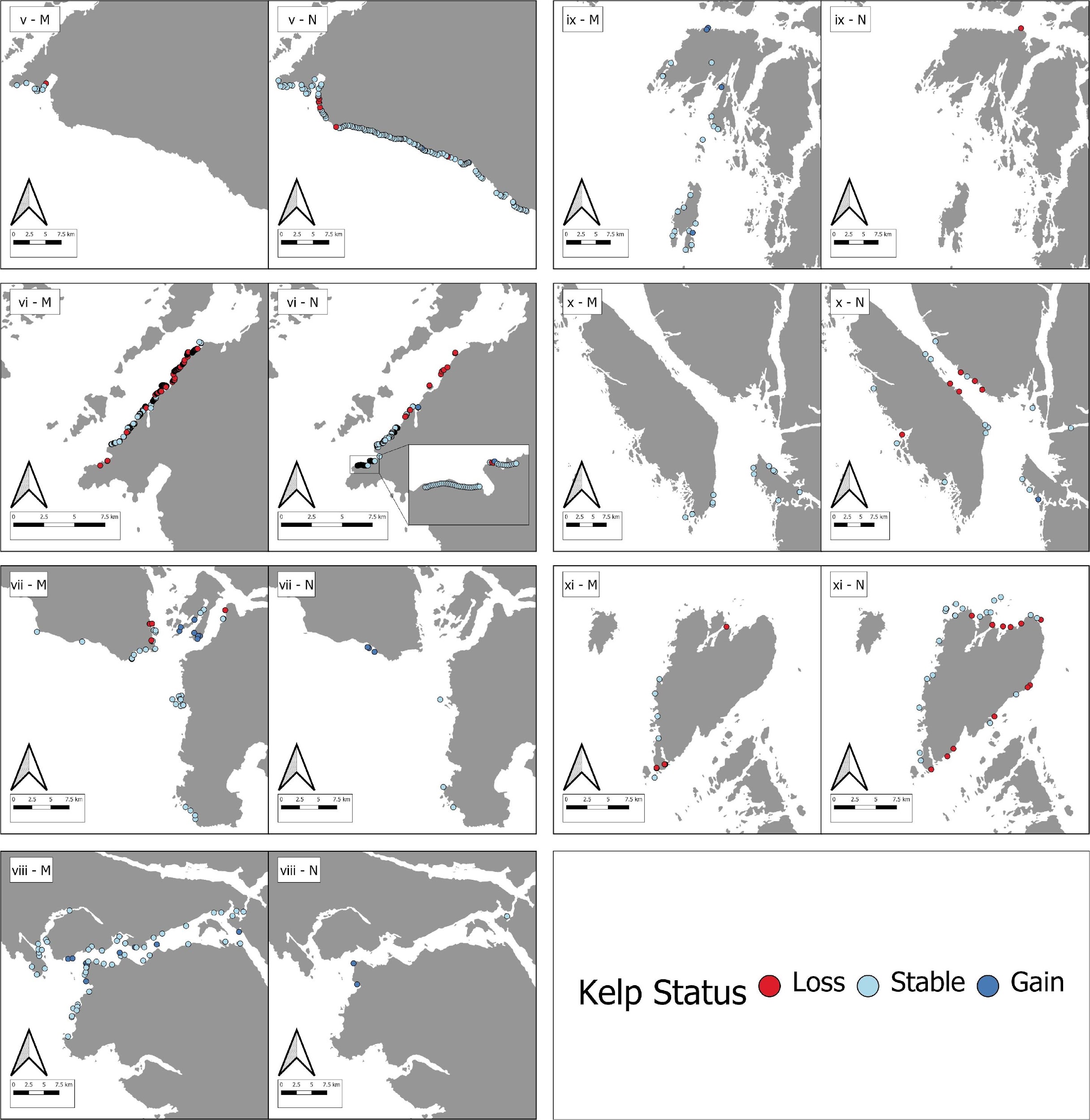

Fig S8. Maps showing species-specific patterns of kelp persistence. Note that points where species could not be discerned were excluded. Labels in the top left of each panel indicate region (v to xi) and species (M = *Macrocystis*, N = *Nereocystis*).

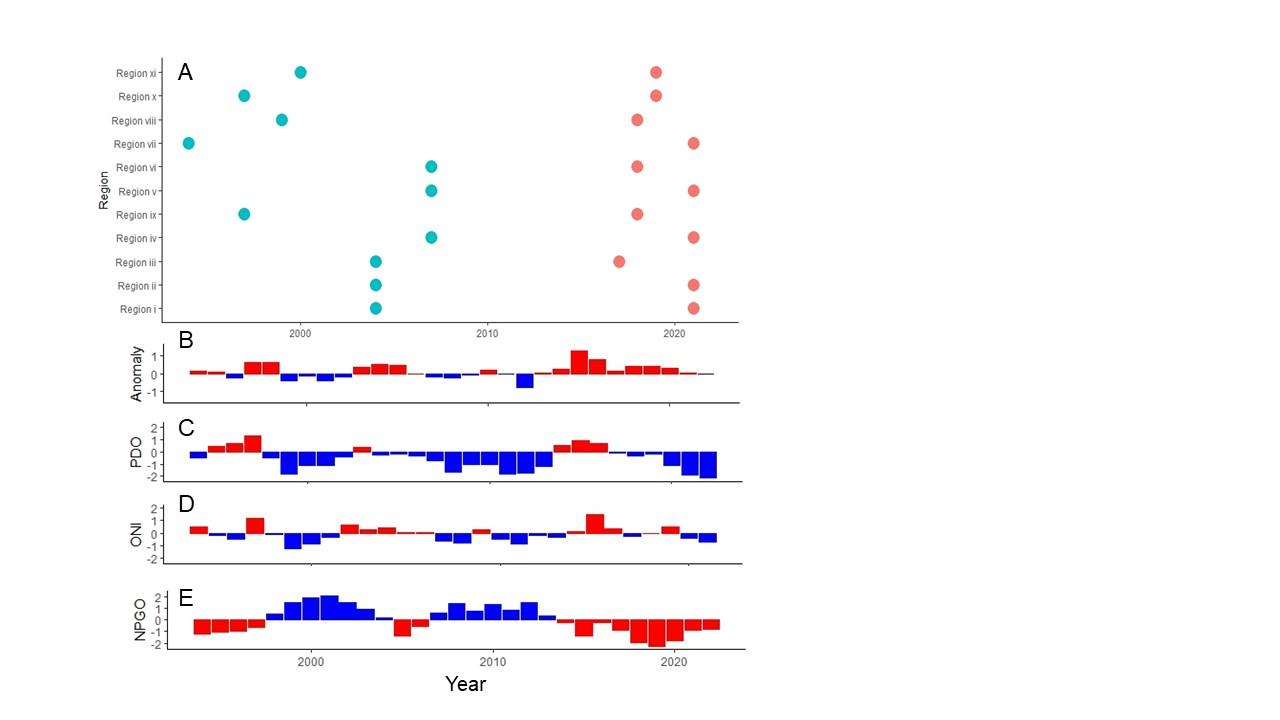

Fig S9. Time points used in snapshot analyses relative to temperature anomalies and climatic oscillators. Shown are the time points from before (teal) and after (red) the heatwave for the snapshot analyses (A). Below are annual mean monthly temperature anomalies (B) and various climate oscillators (C to E). For panels B-E, red colours indicate a warm year and blue represents a cool year. Temperature anomalies represent an average of four lighthouses that span the latitudinal gradient of British Columbia: Amphitrite Point, Chrome Island, McInnes Island and Bonilla Island. Data sources:

PDO: https://www.ncei.noaa.gov/pub/data/cmb/ersst/v5/index/ersst.v5.pdo.dat; ONI: <https://origin.cpc.ncep.noaa.gov/products/analysis_monitoring/ensostuff/ONI_v5.php>; NPGO: http://www.o3d.org/npgo/npgo.php
